# Late-life short-term dietary restriction ameliorates intestinal stem cell function and impacts intestinal stem cell DNA methylation

**DOI:** 10.1101/2024.10.01.616110

**Authors:** Kavitha Kurup, Michael Chan, Eric Moore, Michelle Ranjo-Bishop, Kevin Pham, David R Stanford, Willard M Freeman, Archana Unnikrishnan

## Abstract

Aging reduces the function of intestinal stem cells (ISCs) and dietary restriction (DR) has been shown to improve ISC function in young mice. However, there are no studies to-date that has evaluated the effect of short-term DR (stDR) implemented late-in-life on ISC function. Data from our laboratory shows for the *first time* that *stDR can ameliorate the age-related decline in ISC function by in-vitro* enteroid forming assay. Transcriptomic analysis of the Lgr5^+^ ISCs identified genes not commonly known for ISC function but showed a robust change with age which were reversed by stDR. Evaluation of the methylation status of selected genes (*Slc28a3, Ly9, Cd2, Platr4*) showed association between changes in 5mCG levels in the cis-acting regions of the genes and their expression. Our data suggest a potential role for DNA methylation in stDR’s effect on ISC function.

## Introduction

The stem cell theory of aging posits that age reduces the ability of stem cells to regenerate and replenish the tissues with differentiated cells, leading to decline in the function of the tissue (Chen & Kerr, 2019; Oh et al., 2014). The intestinal epithelium is a stem cell driven tissue that regenerates every 3-5 days throughout adult life which changes in architecture and composition with age due to the declining regenerative capacity of the slowly dividing intestinal stem cells (ISCs) (Nalapareddy et al., 2017; Igarashi & Guarente., 2017). The decline in intestinal regeneration and integrity has been shown to lead to age-related intestinal disorders including intestinal cancers, which are one of the leading causes of death (Nalapareddy et al., 2022; Dai et al., 2022). Therefore, developing strategies to preserve ISC function with age is important to maintaining gastro-intestinal (GI) health and diseases related to GI health, such as insulin resistance (Hseih etal., 2008), cardiovascular disease (Lewis & Taylor, 2020) and neurodegenerative disease (Loh et al., 2024).

DR, the most robust intervention known to delay aging, results in increased lifespan and healthspan and reduced the incidence of most age-related diseases (as reviewed in Green et al., 2022). DR has also been shown to improve the age-related loss in intestinal barrier integrity and reduce the risk of many intestinal disorders, e.g., inflammatory bowel disease, colon cancer, etc. (Akagi et al., 2018; Dai et al., 2022; Rangan et al., 2019; Olivo-Marston et al., 2014). In addition, recent work from our laboratory showed that DR changes the microbial flora of the intestine in aged mice to maintain a more youthful microbiome profile (Kurup et al., 2021). Studies from Yilmaz et al. (2012) & Igarashi & Guarente (2016) have shown that DR increases the number and regenerative capacity of the ISCs through a specialized daughter cell, the paneth cells. Although these studies established the effects of DR on ISC function, these data were from young mice, leaving the effect of DR on ISCs from old mice unexplored and the mechanism of how DR improves ISC function poorly understood.

Emerging evidence shows that DR mediates many of its beneficial effects through an epigenetic mechanism, e.g., DNA methylation, which in turn can affect gene expression over time. Studies on epigenetic clock, the quantitative biomarker of aging developed to predict the age of the animal, have shown that DR slows the epigenetic age of rodent models providing strong evidence that alterations in DNA methylation track with aging (Petkovich et al., 2017; Maegawa et al., 2017; Levine et al., 2020). In addition, several studies including ours have shown that DR alters genome wide CG methylation in various tissues such as liver, brain and kidney (Hahn et al., 2017; Cole et al., 2017; Hadad et al., 2018; Kim et al. 2016a).

In line with the notion that DNA methylation could be mediating DR’s effect, DNA methylation marks (5mC) have been shown to be critical for maintenance of adult stem cells (Chen & Kerr, 2019). Several studies have reported that deficiency in the DNA methylation/demethylation enzymes affects the proliferation and differentiation of adult stem cells, e.g., neural stem cells (Gontier et al., 2018) and ISCs (Kaaij et al., 2013; Sheaffer et al., 2014; Kim et al., 2016b). In addition, Sheaffer et al., (2014) have shown that ISCs have increased expression of Dnmt3a and Dnmt1 suggesting that DNA methylation both de novo and maintenance methylation respectively might be playing a role in ISC proliferation and differentiation. Thus, there is a large body of evidence showing that DNA methylation could be an important mechanism by which DR’s effect on the ISCs is mediated.

In this study, we report for the first time that short-term DR (stDR) implemented late-in-life increases the enteroid forming capacity of ISCs in old mice. We also show that stDR reverses the expression of ISC genes found to change with age and that these changes in expression are associate with changes in DNA methylation in the cis-acting regions of the gene.

## Results

### Age reduces enteroid forming capacity of ISCs

Nalapareddy et al. (2017) have shown that intestinal crypts from 22-month-old mice show reduced regenerative capacity *in vitro*, therefore, we first tested the potential of crypts isolated from 6- and 25-month-old mice to form enteroids. As previously shown, intestinal crypts that include stem cells have the ability to regenerate *in vitro* into clonal, multipotent enteroids with all specialized intestinal cells (Griegroff et al., 2015). Figure 1A shows images of multilobed enteroids formed on day 5 from crypts and Figure 1B quantitatively shows that age reduced by 50% the total number of enteroids formed from crypts of old mice compared to the young mice.

**Figure 1:**
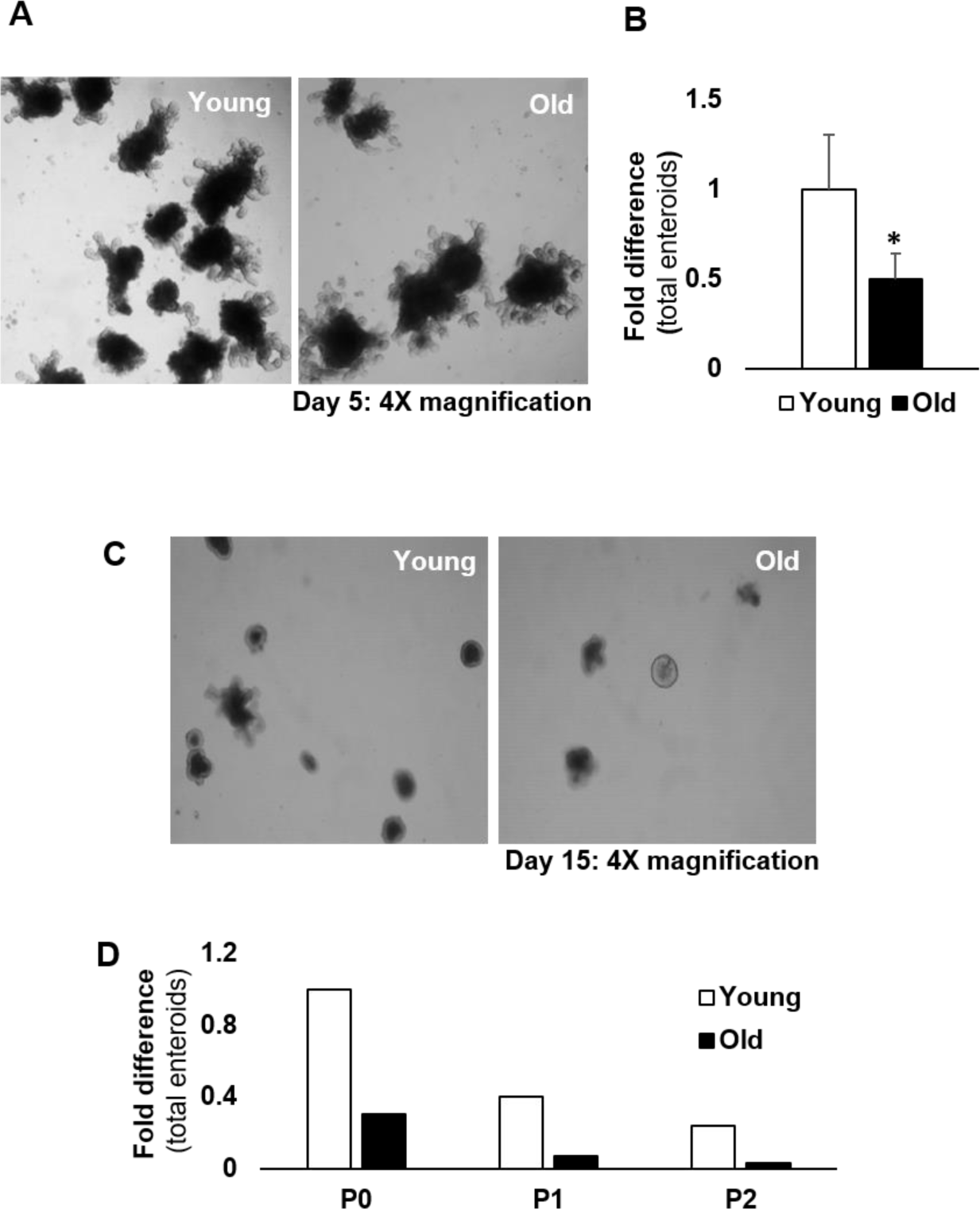
Enteroid formation in Young and old male mice. (A) Enteroids formed from intestinal crypts. 4X representative images of enteroids at day 5 in culture is shown. (B) fold difference of the total enteroids formed. The data shown are the mean ± SEM from 6 mice/group at 6- and 25-months of age. *<0.05 with student’s t-test. (C) Enteroids formed from freshly isolated Lgr5^+^ ISCs isolated from young and Old male mice. 4X images of enteroids at day 15 in culture is shown. The enteroid images shown are representative images from 6 mice/group. (D) Fold difference of total enteroids formed from freshly isolated Lgr5^+^ ISCs when they are passaged (P0-first enteroids formed from fresh ISCs; P1-enteroids formed after one passage of enteroids from P0; P2-enteroids formed after two passages of enteroids from P0). The fold difference in enteroids shown for passaging data is from 2 mice/group; About 12 replicates were set up for each mice/group and enteroids from each well followed for passaging.

Next, we tested the effect of age on the regenerative capacity of ISCs (without the stem cell niche) using ISCs marked by the presence of leucine-rich repeat containing G-protein coupled receptor 5 (Lgr5^+^ ISCs). Lgr5^+^ ISCs are small cycling ISCs located at the intestinal crypt base and have the ability to differentiate into other epithelial lineages (Yan and Kuo., 2015), including enteroids that are indicative of a normal intestine in culture (Sato et al., (2009). For this experiment we isolated Lgr5^+^ ISCs from young (6-month) and old (25-month) mouse intestinal crypts with anti-Lgr5^+^ antibodies conjugated to magnetic beads. Magnetic bead sorting provides cell isolation with high purity, viability and minimal stress to the cells such that the cells can be used efficiently for downstream analysis such as cell culturing. S1A shows the mRNA expression of ISC markers (Lgr5^+^, Olfm4, Ascl2) confirming that our bead sorted ISC population consisted of Lgr5^+^ ISC population. To further confirm that the anti-Lgr5^+^ antibody was specific for the Lgr5^+^ ISC population and not for paneth cells that are found in close proximity to the ISCs in the intestinal crypt, we measured the expression levels of paneth cell markers (Defa 22, Lyz1) in the ISC pool sorted using anti-Lgr5^+^ antibodies and compared it to the levels of these markers in paneth cell population sorted using anti-CD24^+^ antibodies (S1B). As can be seen in S1B, Lgr5^+^ ISC pool had a low-level expression of paneth markers compared to the CD24^+^ paneth cell pool showing that our Lgr5^+^ ISC pool is predominantly made up of Lgr5^+^ ISCs.

We next evaluated the enteroid forming capacity of the freshly isolated Lgr5^+^ ISCs. Sato et al. (2009) showed that a single Lgr5^+^ ISC from young (∼3-months) mice can form a multilobed enteroid by day 13. Lgr5^+^ ISCs isolated from 6-month-old differentiated into multilobed enteroids by day 15 as described in Sato et al., (2009). In contrast, Lgr5^+^ ISCs isolated from old mice showed reduced formation of multilobed enteroids even by day 15 compared (Figure 1C). Some ISCs from old mice, which started developing into enteroids normally (about 60%), progressed until day 5 but did not develop into multilobed enteroids. Next, we took the enteroids formed from either Lgr5^+^ ISCs isolated from young or old mice, dissociated them into single cells and replated the single cells to form new enteroids. We did this passage twice (P1 and P2), and Figure 1D shows that the Lgr5^+^ ISCs isolated from young mice formed viable multilobed enteroids after P1 and P2, whereas the Lgr5^+^ ISCs isolated from old mice showed reduced enteroid forming capacity on passaging and by the second passage (P2) no viable enteroids were observed confirming that aging reduces the enteroid forming capacity of the ISCs in culture.

### Short-term dietary restriction (stDR) enhances the enteroid forming capacity of ISCs in both young and old

To determine the effect of stDR on the regenerative capacity of Lgr5^+^ ISCs, we fed 2-and 21-months old mice 40% DR for 4-months and compared them to mice fed *ad-libitum* (AL). stDR reduced body weight in both young and old by 36% and 24% respectively (S2A) and reduced gonadal fat (>75%) in young and old mice (S2B) similar to our previously published work (Matyi et al., 2018). We first evaluated the enteroid forming capacity of the Lgr5^+^ ISCs isolated from young mice fed DR diet for 4-months. stDR increased the formation of total enteroids from the young ISCs by ∼11-fold (Figure 2A&B) and also increased the number of multilobed enteroids (>2 lobes) by ∼10-fold compared to their AL counterparts (Figure 2C).

**Figure 2:**
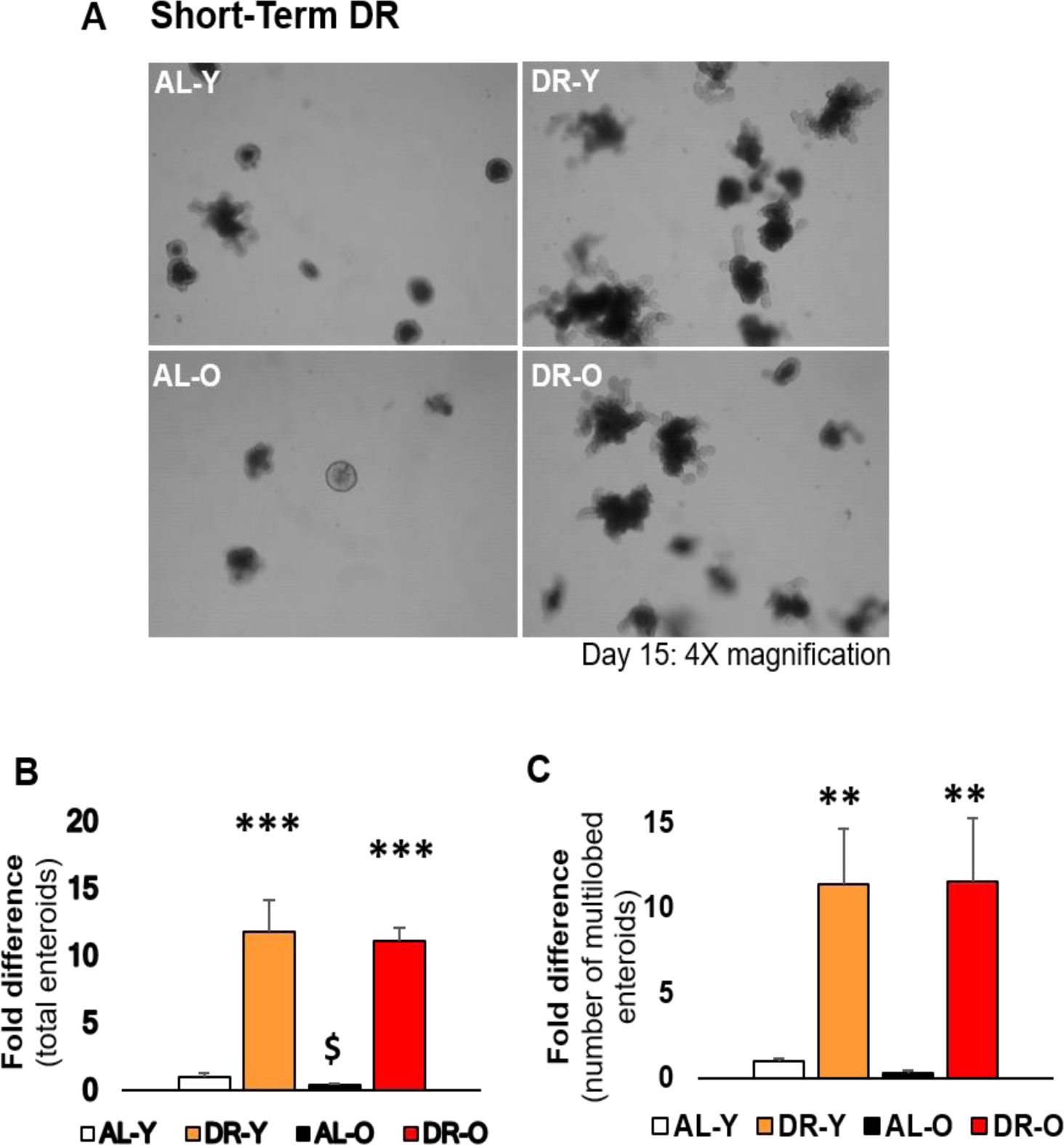
Enteroid formation from freshly isolated Lgr5^+^ ISCs isolated from young (Y) and old (O) AL and DR male mice fed short-term DR for 4 months. (A) 4X images of enteroids at day 15 in culture. (B) fold difference of the total enteroids formed (C) fold difference of the number of multilobed enteroids (enteroids with more than 2 lobes). The data shown are the mean ± SEM from 6 mice/group. ***<0.0001 with one-way ANOVA and Tukey’s multiple comparison test. All means compared to mean of AL-Y. $p<0.05 with two-tailed students T-test comparing AL-O to AL-Y.

Figure 2A & B (bottom panel) shows the enteroid forming capacity of the Lgr5^+^ ISCs isolated from old mice fed stDR diet for 4-months. The enteroid forming capacity of the ISCs obtained from old mice was reduced ∼50% compared to young mice. However, stDR increased enteroid forming capacity ∼11-fold, which was to the same extent as observed in the young ISCs exposed to stDR. In addition, stDR also increased the number of multilobed enteroids (>2 lobes) by ∼10-fold by day 15 in the old mice (Figure 2C). Taken together, our data show for the *first time* that stDR can rescue the age-related decline in ISC enteroid forming capacity.

To characterize the enteroids formed from Lgr5^+^ ISCs obtained young and old AL and stDR mice we determined the expression of key genes involved in cellular regeneration (Wnt10b, Tert), quiescence (Lrig1, Bmi1) and cell cycle arrest (Cyp2b10lp16). For this, we used data from RNA-seq analysis performed only on the multilobed enteroids found at day 15. As shown in S3, we found that stDR reduced the expression of quiescence and cell cycle arrest markers and increased the expression of regeneration markers in enteroids from both young and old but more robust in the old mice (S3A, B, C). These key gene expression changes in the enteroids show that stDR could be increasing the regenerating capacity of the enteroids by inducing expression of genes that promote regeneration.

### Age alters and stDR reverses the transcriptomic changes in the ISCs

To gain insight into the potential mechanism underlying stDR’ s effect on ISC regeneration in culture, we studied the effect of age and stDR on the transcriptome of Lgr5^+^ ISCs freshly isolated from young and old mice fed AL or DR for 4-months. Differential expression analysis showed that the expression of 1338 ISC genes changed significantly [p (Corr) < 0.05 and 2-fold change] with age (old AL vs young AL) and stDR (Old DR vs old AL and young DR vs young AL). PCA by sample profiles of differentially expressed genes in young and old mice fed AL and stDR demonstrated that the old AL clearly separated from the young AL whereas both the young and old stDR clustered near the young AL. While the young stDR clustered closer to the young AL, the old stDR was clustered away from the old AL, (Figure3A) showing that age shifted the transcriptomic profile and stDR in the old shifts them back closer to the young.

Of the 1338 differentially expressed ISC genes, 895 changed with age (447 increased and 448 decreased in expression in the Lgr5^+^ ISCs isolated from young and old mice), and 678 (75%) of these genes were reversed by stDR (408 reduced and 270 increased) in the Lgr5^+^ ISCs obtained from old mice (Figure 3B; S4A&B). stDR also significantly changed the expression of 131 genes independent of age with 124 genes showing reduced and 7 genes showing increased expression in the Lgr5^+^ ISCs from old mice (Figure 3B; S4A&B). Further, stDR significantly altered the expression of 414 genes in young mice (164 down and 250 up with DR) (Figure 3B; S4A&B). A total of 38 genes were found to be common between all four conditions studied (Figure 3B & C). None of the 38 genes identified are genes traditionally known to be involved in ISC function. A major shift in these 38 genes occurs in Lgr5^+^ ISCs from old AL mice compared to Lgr5^+^ ISCs from young AL mice. Interestingly, stDR shifted the expression of these 38 in a similar direction for Lgr5^+^ ISCs isolated from either young or old mice, suggesting that these 38 genes could play an important role in the mechanism responsible for the ability of stDRs to increase the regenerative potential of Lgr5^+^ ISCs isolated from both young and old mice.

**Figure 3:**
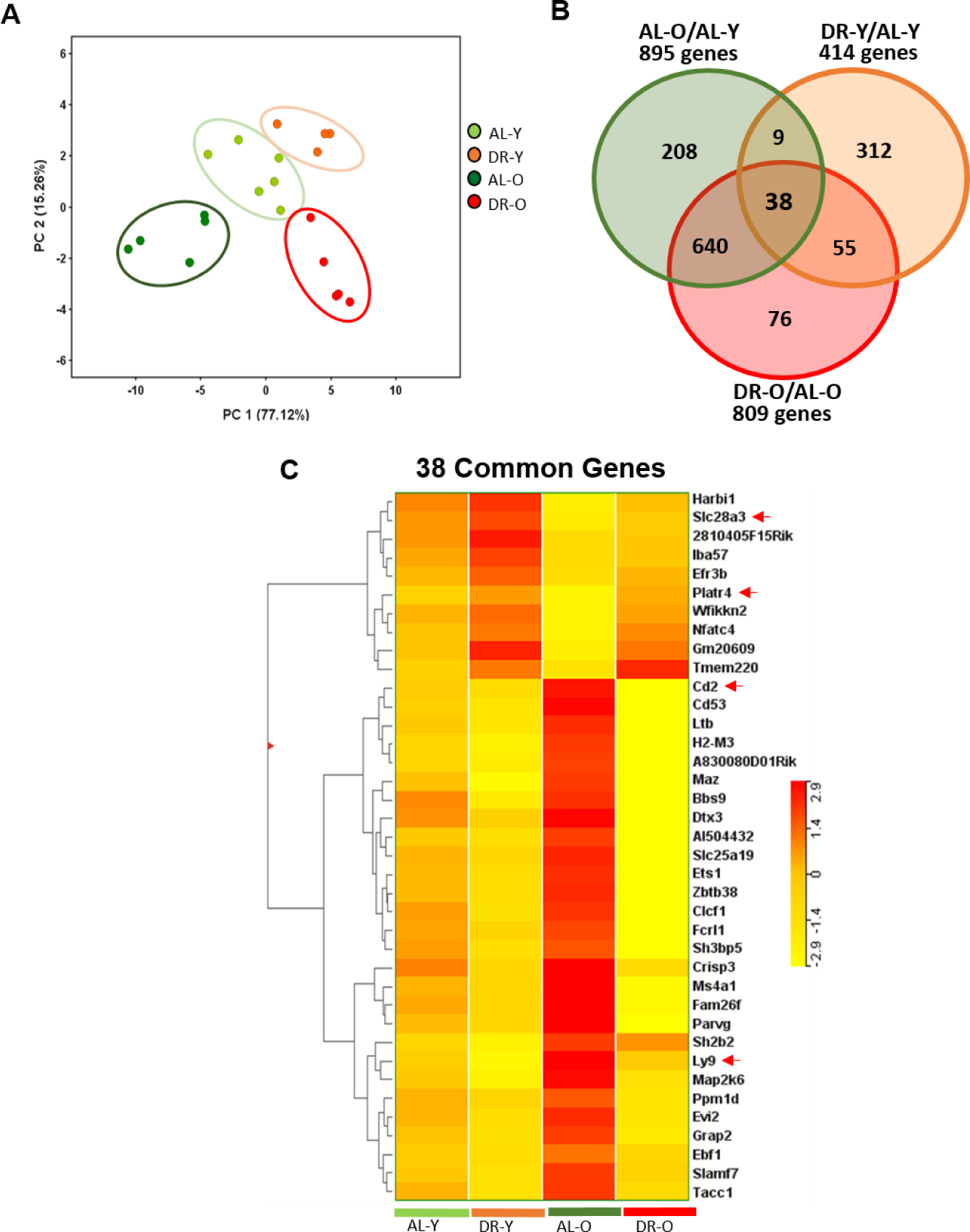
Comparison of the transcriptome of ISCs from young (Y) and old (O) male mice fed AL or stDR for 4-months. (A) PCA plots showing 2-fold change generated using data of normalized signal values with one-way ANOVA p cut-off 0.5 and FC cut-off 2. (B) Venn diagram with the total number of transcripts that increased or decreased with age and stDR. (C) heat map of the relative levels of the 38 transcripts found to be common between all four groups studied that showed significant change using one-way ANOVA with Benjamini Hotchberg correction at p ≤ 0.05 and fold change ≥ 2. The data are the mean ± SEM from 4-6 mice/group (6 (AL-Y), 4 (DR-Y), 5 (AL-O) and 5 (DR-O)) at 6- and 25-months of ages fed AL or 40% DR for 4 months. DR was started at 2-months and 21-months of age.

### Short-term DR alters DNA methylation of candidate genes in the CG context

To investigate DNA methylation as the regulatory mechanism underlying DR’s effect on the transcriptome, we measured DNA methylation using oxidative whole genome bisulfite sequencing (oxWGBS), which allowed us to accurately measure the methylated cytosines (5mC’s) because oxidative bisulfite sequencing differentiates between 5Cs and 5hmCs (Booth et al., 2013). In this series of experiments, we first measured the global 5mCs in the CG context in the Lgr5^+^ ISCs freshly isolated from young and old mice fed either AL or the stDR diet (S5). We focused on DNA methylation changes in the CG context because our previous study showed that stDR changes methylation level in the CG sites in the intestinal mucosa (Unnikrishnan et al., 2017). Moreover, DNA methylation changes in CG sites found in ISC proliferation and differentiation genes correlate to gene expression (Sheaffer et al., 2014) making this an important DNA modification to study. As shown in S5, although stDR reduced 5mCG count in ISCs from young mice it was not significant. However, in old mice, a 6% decrease in 5mCG count was observed with stDR that was statistically significant. To determine if DNA methylation played a role in the expression of genes observed to change with age and stDR, we chose 4 representative genes (*Slc28a3, Ly9, Cd2, Platr4*) to study from the 38 genes found to be common between four groups with their expression pattern changing in opposite direction with age and DR. For example, *Slc28a3* and *Platr4* expression decreased with age and stDR increased their expression while, *Ly9* and *Cd2* expression increased with age and stDR reduced it. In these experiments, the total 5mCG levels were measured in the 4000bp-nucleotide region upstream and downstream of the transcription start site (TSS) of each candidate gene that would cover both cis- and trans-acting regulatory regions of the gene. And within the −4000 to +4000 region, we focused our analysis on the cis-acting region (−500 to +500 around TSS) which has been traditionally shown to regulate gene expression (Bird, 2002).

First, we evaluated the expression and methylation status of *Slc28a3* (solute carrier family 28-member 3), which is a nucleoside transporter known to be involved in nucleoside uptake in various cell types including stem cells (Errasti-Murugarren et al., 2009; Anabtawi et al., 2023; Hesse et al., 2024). Figure 4A shows that mRNA levels of *Slc28a3* decreased ∼2.5-fold in Lgr5^+^ ISCs isolated from old mice compared to young mice. The expression of *Slc28a3* was increased ∼1.5 fold by stDR for Lgr5^+^ ISCs isolated from both young and old mice. The 5mCG levels of the 4000bp-nucleotide region upstream and downstream of the TSS, the gene body and the −500 to +500 cis-acting region of the *Slc28a3* from Lgr5^+^ ISCs are shown in S6. None of these regions showed a significant change in total mCG levels with either age or stDR. However, we identified a 100bp region (+200 to +300) on the 3’ flanking side of the TSS of the *Slc28a3* gene which showed DNA methylation changes in the Lgr5^+^ ISCs that were associated with the changes in the expression of the gene (Figure 4B). The 100bp region which encompasses the first intronic region of the *Slc28a3* gene body has 3 CG sites (Figure 4C), and the total methylation at these 3 CG sites showed a modest increase with age, which was not significance; however, the total 5mCG levels in this region were significantly reduced (>85%) by stDR in both the young and old mice compared to their AL counterparts (Figure 4B). Thus, hypomethylation in this region of the *Slc28a3* gene by stDR is associated with the increased mRNA expression. Studies have consistently shown that methylation in the first intronic region has an inverse relation with gene expression (Anastasiadi et al., 2018), supporting the association we found between the methylation and gene expression of Slc28a3.

**Figure 4:**
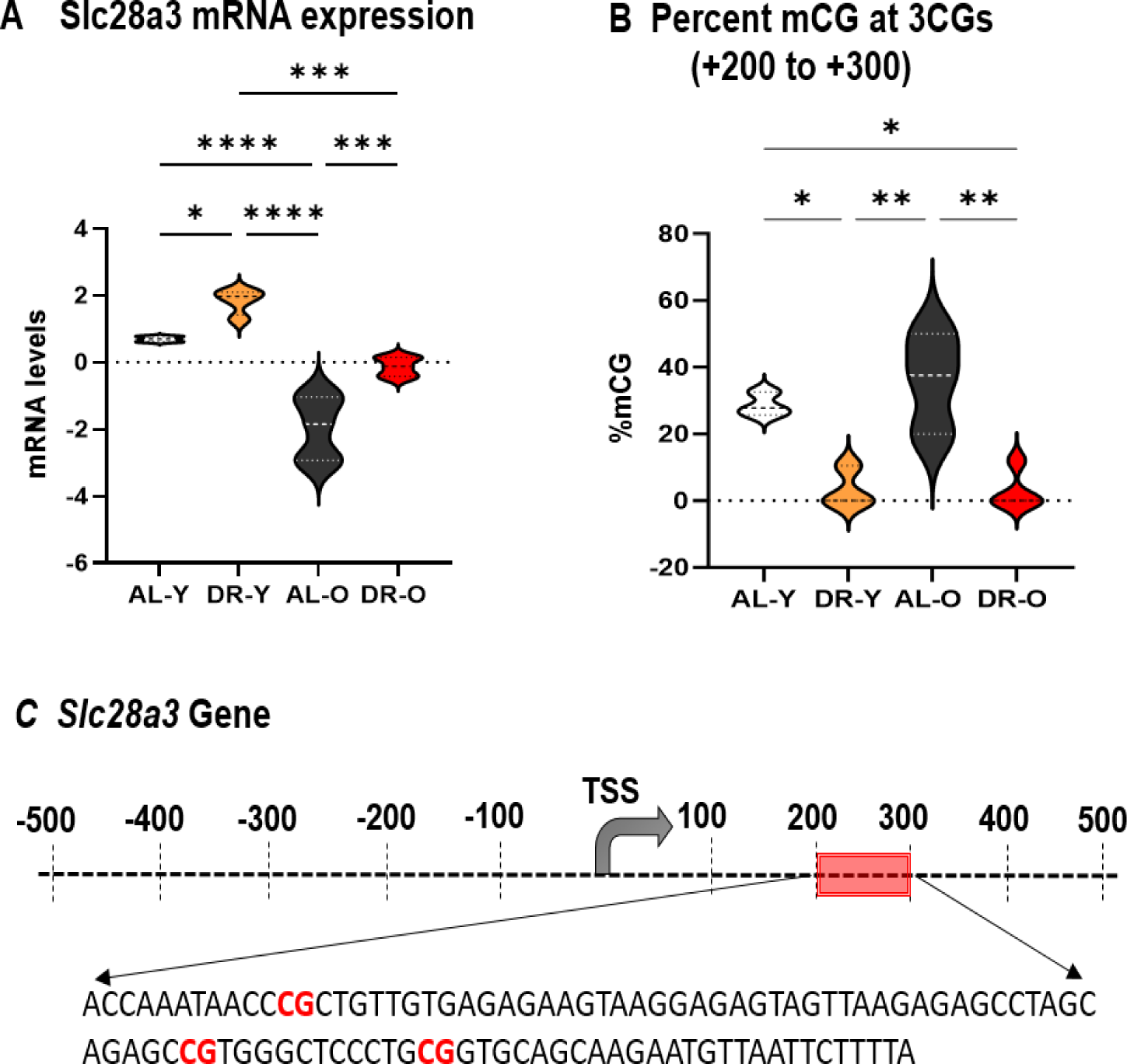
Gene expression and DNA methylation profile of *Slc28a3* gene in young (Y) and old (O) mice fed AL or 40% DR for 4-months. (A) mRNA levels of *Slc28a3* gene from the RNA-seq data. (B) total percent methylation (%mCG) of 3CGs in the +200 to +300 region downstream of TSS encompassing first intron. (C) Line map of the −500bp to +500bp region of the *Slc28a3* gene around the TSS with the sequence of a 100bp differentially methylated intron region. The data is from Lgr5^+^ ISCs freshly isolated from 6- and 25-month-old C57BL/6 male mice fed AL or 40% DR for 4-months. The mRNA data (from RNA-Seq) shown are the mean ± SEM from 4-6 mice/group (6 AL-Y, 4 DR-Y, 5 AL-O, 5 DR-O). The %mCG data are the mean ± SEM from 3-4 mice/group. DR was started at 2-months and 21-months of age. The violin plots show median and the quartiles. *p<0.05, ** p<0.01, *** p<0.001, ****p<0.0001 with one-way ANOVA with tukey’s multiple testing correction.

We next studied *Ly9* (*CD229*) in the Lgr5^+^ ISCs that is a cell-surface receptor, found in certain subsets of adult stem cells (Somuncular et al., 2023; Yamada et al., 2015). Expression of Ly9 mRNA was increased ∼3-fold with age (Figure 5A) and stDR reduced Ly9 expression in Lgr5^+^ ISCs isolated from both young (∼2-fold) and old mice (3-fold) when compared to their AL counterparts. As shown in S7A, B, C, there was no difference in the percent mCG methylation in the 4000bp region upstream and downstream of the TSS and in the gene body region. However, stDR was found to significantly increase CG methylation in the −500 to +500bp region around the TSS by ∼10-fold in ISCs from young mice and by 2-fold in old mice compared to their AL counterpart (S7D). Within the −500 to +500bp region we identified a 100bp site (−100 to +1) upstream of the TSS containing 3CG sites (Figure 5C) that showed a difference in methylation. Total mCG levels in Lgr5^+^ ISCs did not change significantly with age. However, stDR led to a ∼70-fold increase in total mCG levels in Lgr5^+^ ISCs from young mice and ∼4-fold increase in old mice (Figure B). The increased methylation with stDNA was associated with a significant decrease in Ly9 expression by stDR in Lgr5^+^ ISCs from old mice. In young mice, the increase in methylation in ISC was observed with stDNA was also associated with a >80% decrease in Ly9 expression; however, this decrease was not statistically significant.

**Figure 5:**
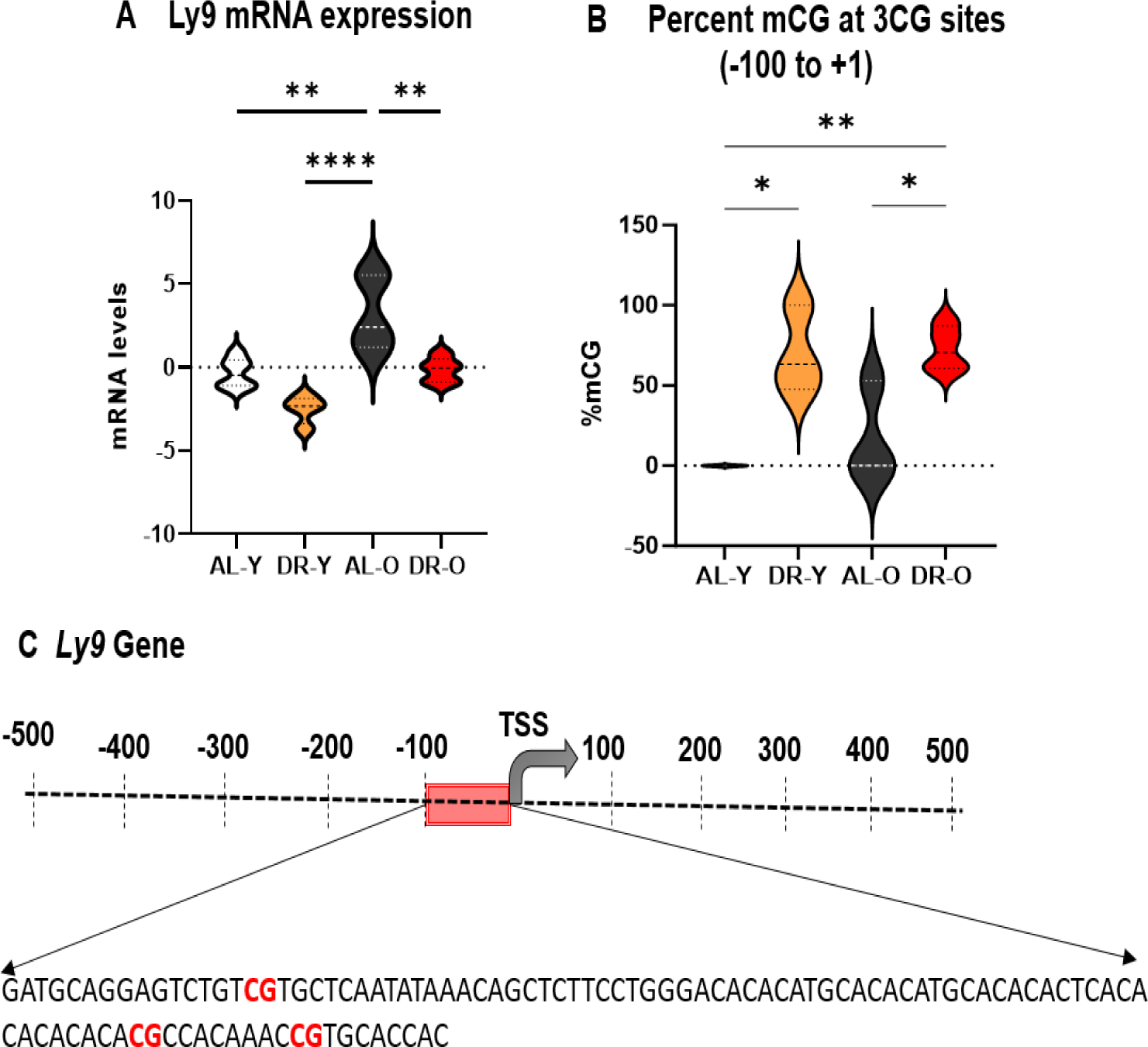
Gene expression and DNA methylation profile of *Ly9* gene in young (Y) and old (O) mice fed AL or 40% DR for 4-months. (A) mRNA levels of *Ly9* gene from the RNA-seq data. (B) total percent methylation (%mCG) of 3CGs in the 100bp region (−100 to +1) upstream of the TSS. (C) Line map of the −500bp to +500bp region of the *Ly9* gene around the TSS with the sequence of the 100bp differentially methylated region. The data is from Lgr5^+^ ISCs freshly isolated from 6- and 25-month-old C57BL/6 male mice fed AL or 40% DR for 4-months. The mRNA data (from RNA-Seq) shown are the mean ± SEM from 4-6 mice/group (6 AL-Y, 4 DR-Y, 5 AL-O, 5 DR-O). The %mCG data are the mean ± SEM from 3-4 mice/group. DR was started at 2-months and 21-months of age. The violin plots show median and the quartiles. *p<0.05, ** p<0.01, ****p<0.0001 with one-way ANOVA with tukey’s multiple testing correction.

The third gene we studied was Cd2, a cell adhesion molecule shown to be expressed in spermatogonial stem cells in mice and rats (Espagnolle et al., 2007; Kanatsu-Shinohara et al., 2020). The expression of Cd2 mRNA was increased ∼2.5-fold in Lgr5^+^ ISCs from old mice compared to young mice (Figure 6A), which was significant with fisher’s LSD test. stDR did not alter Cd2 mRNA expression in young mice but significantly reduced (∼7.5 fold) Cd2 mRNA expression in old mice compared to old AL mice. Next, we analyzed the methylation status of the 4000bp region upstream of the TSS of the Cd2 gene. Age increased the total percent mCG levels by ∼1.5 fold (S8A). stDR also significantly increased the mCG levels in the young and old compared to the young AL (S8A). As shown in S8B the total mCG levels did not significantly differ in the 4000bp downstream region. Age increased the gene body methylation by ∼1.2 fold and stDR increased the gene body methylation in the young and old compared to the young AL as well (S8C). Age and stDR seemed to increase overall methylation of the cis-and trans-acting region of the Cd2 gene but when we evaluated the mCG levels of −500 to +500 region around the TSS (S8D), we did not find any difference with age but observed an increasing trend with stDR (significant in stDR old mice). Subsequently we narrowed down our analysis to a 200bp region (−200 to +1) with 3CG sites. Here we found a modest reduction (not significant) in methylation with age but stDR in old mice increased the percent 5mCG methylation ∼ 64-fold compared to old AL mice (Figure 6B). Figure 6C shows the sequence of the −200 to +1 region with the 3CG sites. Although the increase in Cd2 mRNA expression with age was not associated with mCG levels, the decreased mRNA expression observed with stDR in the old mice is associated to increased methylation in the −200 to +1 region studied.

**Figure 6:**
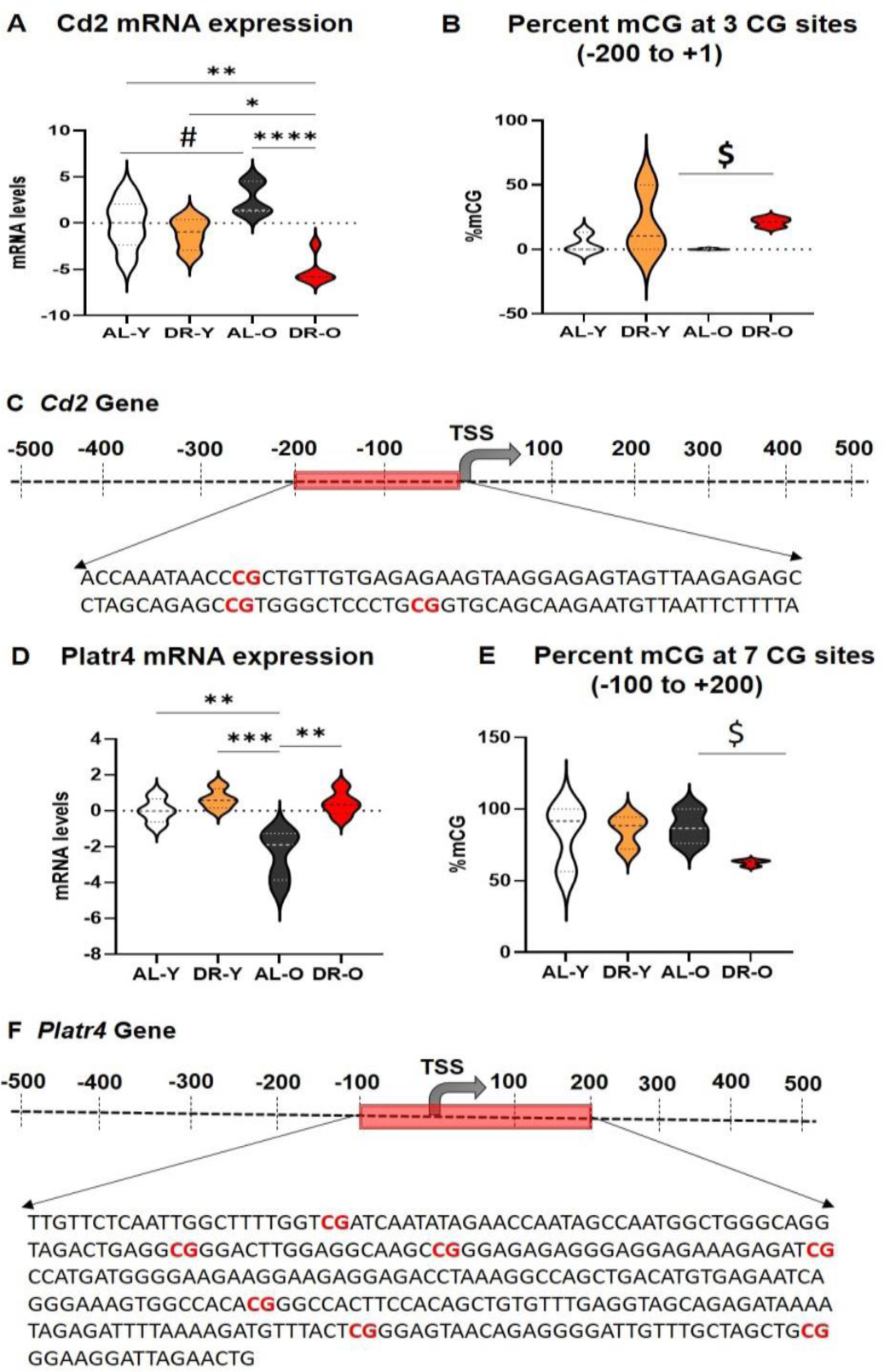
Gene expression and DNA methylation profile of *CD2* and *Pltar4* gene in young (Y) and old (O) mice fed AL or 40% DR for 4-months. (A) mRNA levels of *CD2* gene from the RNA-seq data. (B) total percent methylation (%mCG) of 3CGs in the 200bp region (−200bp to +1bp) upstream of the TSS of CD2 gene. (C) Line map of the −500bp to +500bp region of the *CD2* gene around the TSS with the sequence of the 200bp differentially methylated region. (D) mRNA levels of *Platr4* gene from the RNA-seq data. (E) total percent methylation (%mCG) of 7CGs in the 300bp region (−100 to +200) around the TSS encompassing the first exon and intron. (F) Line map of the −500bp to +500bp region of the *Platr4* gene around the TSS with the sequence of a 300bp region encompassing the first exon and intron. The data is from Lgr5^+^ ISCs freshly isolated from 6- and 25-month-old C57BL/6 male mice fed AL or 40% DR for 4-months. The mRNA data (RNA-Seq) shown are the mean ± SEM from 4-6 mice/group (6 AL-Y, 4 DR-Y, 5 AL-O, 5 DR-O). The %mCG data are the mean ± SEM from 3-4 mice/group. DR was started at 2-months and 21-months of age. The violin plots show median and the quartiles. *p<0.05, ** p<0.01,****p<0.0001 with one-way ANOVA with tukey’s multiple testing correction. #p<0.05 with one-way ANOVA with uncorrected fisher’s LSD test. $p<0.05 with two-tailed students T-test comparing DR-O to AL-O.

The last candidate gene we studied was *Platr4* (pluripotency-associated transcript 4) an early embryonic long non-coding RNA (LncRNA). Platr4 mRNA decreased significantly (2.5-fold) with age in Lgr5^+^ ISCs (Figure 6D). stDR did not alter the mRNA expression in the young mice but significantly increased Platr4 expression (>2-fold) in the old mice (Figure 6D). As shown in S9, total 5mCG levels did not change significantly with age or stDR in Lgr5^+^ ISCs in the 4000bp-nucleotide region upstream and downstream of the TSS, the gene body and the −500 to +500 cis-acting region of the *Platr4* gene. However, in the 300bp region around the TSS (−100 to +200) encompassing the first exon and intron of the gene, which contained 7CG sites (Figure 6F), stDR significantly reduced the mCG levels in the region in Lgr5^+^ ISCs from old mice (Figure 6E). While the decrease in mRNA expression observed with age was not associated with any changes in DNA methylation, the increase in mRNA expression observed with stDR in old mice was associated with a reduction in DNA methylation.

## Discussion

It is well documented that the intestinal regenerative capacity declines with age and is associated with an age-related decline in the proliferative capacity of Lgr5^+^ ISCs (Moorefield et al., 2017; Mihaylova et al., 2018; Nalapareddy et al., 2022; Choi et al., 2023). In addition, several studies have shown that feeding young (≤6 months of age) mice a DR-diet for 4 to 28 weeks increased ISC number and improved the ability of ISCs obtained from the mice to regenerate in culture (Yilmaz et al., 2012; Igarashi & Gaurente 2016). The goal of this study was to determine if 4 months of DR started at 21-months of age could reverse the decreased function of ISCs that others have observed in Lgr5^+^ ISCs isolated from old mice, e.g., the ability of Lgr5^+^ ISCs to regenerate in culture is dramatically reduced in 18-to 22-month-old mice (Nalapareddy et al., 2017; Moorefield et al., 2017; Mihaylova et al., 2018). Our data clearly demonstrate that initiating DR late-in-life can rescue the reduced ability of Lgr5^+^ ISCs from old mice to regenerate and form enteroids. In fact, the function of Lgr5^+^ ISCs isolated from 25-month-old mice fed DR for 4 months was not only rescued but their ability to form enteroids was even greater than ISCs isolated from young, 6-month-old mice. To our knowledge, this is the first study to show that implementing DR late-in-life can reverse a physiological function that has declined significantly in an old animal. It is well documented that gastro-intestinal (GI) integrity and function declines with age and that life-long DR maintains a better GI system (Heller et al., 1990; Holt et al., 1991). Our data are particularly important because it suggests that it might be possible to restore GI function in older individuals through short-term DR or other anti-agin0g strategies that mimic DR such as intermittent fasting and rapamycin.

While our data and previous studies show that DR improves the function of ISCs, the mechanism(s) for how DR alters Lgr5^+^ ISC function is poorly understood. We previously showed that life-long DR prevents more than 75% of transcriptomic changes observed with age in the intestinal mucosa (Kurup et al., 2021). Ironically, there are no reports that have studied the impact of age and stDR on the Lgr5^+^ ISC transcriptome. Therefore, we compared the transcriptome of Lgr5^+^ ISCs isolated from young and old mice fed either AL or a stDR diet. Age had a dramatic impact on the transcriptome of the Lgr5^+^ ISCs, e.g., the transcripts of 895 genes were significantly changed (>=2.0-fold) with age, and stDR reversed the changes in ∼ 75% (678 genes) of the transcripts that changed significantly in the old mice fed AL. Of all the genes that changed significantly with age and stDR, we found only 38 genes to be common between the four groups we studied (young and old mice fed AL and stDR). These genes, although not associated with ISC function to-date, were found to change significantly by ≥2-fold with age, suggesting they could play an important role in the decline of ISC function with age. On further analysis using GO function (data shown in supplement: S4C), we found that these 38 genes had predominant roles in cellular/biological functions as cell surface markers, immune response and membrane protein binding functions.

Once we found that stDR could attenuate the age-related changes in the expression of the majority of genes that changed with age, we were interested in exploring the mechanism of how DR might alter the expression of genes in ISCs. We rationalized that DNA methylation could play a role in the changes in the transcriptome observed with stDR in Lgr5^+^ ISCs for the following reasons. *First*, there is a large body of evidence showing that DNA methylation regulates the expression of various genes involved in stem cell proliferation and differentiation (Sheaffer et al., 2014, Kim et al., 2016b, Gontier et al., 2018). For example, Kaaij et al., 2013 showed that genes with hypomethylation in the enhancer regions showed increased expression during ISC differentiation. Further, Sheaffer et al., 2014 reported that a deficiency of Dnmt1 lead to altered methylation in ISC genes (e.g., Wnt responsive genes), which in turn led to dysregulated gene expression and delayed differentiation of ISCs. *Second*, DR has been shown to alter DNA methylation in various tissues (for review see Zhai et al., (2023). For example, genome wide CG methylation was altered by DR in liver (Hahn et al., 2017; Cole et al., 2017), and we showed that 35 to 38% of the age-related methylation changes observed in the hippocampus of 24-month-old male fed AL were prevented by DR (Hadad et al., 2018). Kim et al. (2016a) also reported that 4-weeks of DR in old rats altered whole genome DNA methylation in kidney. *Third*, we have shown that age reduces, and DR restores the expression of genes involved in DNA methylation e.g., Dnmts and Tets in the intestinal mucosa (Unnikrishnan et al., 2018). In addition, we found that DR implemented for 4-months in young mice decreased methylation in the 397bp promoter region of the Nts1 gene (with three CG sites) and concomitantly increased the expression of Nts1 mRNA in intestinal mucosa (Unnikrishnan et al., 2017).

To study the role of DNA methylation in the changes in gene expression we observed in the Lgr5^+^ ISCs, we chose four representative genes (*Slc28a3, Ly9, Cd2, Platr4*) from the 38 common genes because they changed significantly (2-fold) with age and the age-related changes were reversed by stDR in old mice. Using oxWGBS, we were able to accurately measure true mCG levels in the −4000 to +4000bp region surrounding the TSS of each of the four genes. We found that stDR significantly altered methylation in the cis-acting region −500 to +500 around the TSS and these changes were associated with changes in the expression of genes induced by stDR in the Lgr5^+^ ISCs of old mice. For example, the increased expression of *Slc28a3* and *Platr4* induced by stDR in old mice was associated with decreased methylation. On the other hand, the decreased expression of *Ly9* and *Cd2* induced by stDR in old mice was associated with increased methylation. We found similar associations between methylation and expression in *Slc28a3* and *Ly9* by stDR in young mice. However, expression and methylation of *Cd2* and *Platr4* showed modest or no significant changes with stDR in ISCs isolated from young mice.

Previous reports have shown that expression of *Slc28a3* and *Cd2* was correlated to DNA methylation. Hua et al., (2022) showed that *Slc28a3* gene is regulated by DNA methylation in mothers at advanced age during pregnancy (AMA-advanced maternal age) and her offspring. The AMA mother and offspring had reduced Slc28a3 gene expression, which was associated with an increased DNA methylation pattern especially in a differentially methylated region, located in the intron 1 of the gene. Interestingly, we also found DNA methylation in the intron 1 region of *Slc28a3* to be significantly reduced by stDR in Lgr5^+^ ISCs and the decreased methylation was associated with an increase in the expression of Slc28a3 mRNA. Similarly, methylation at CG sites within the promoter region and exon 1 of the Cd2 gene was reported to regulate the expression of Cd2 in T-lineage tumor cell lines (Wotton et al., 1989). In Lgr5^+^ ISCs, we found the region 500bp upstream and downstream of the TSS of Cd2, which encompasses the promoter region and the exon1 (including data shown in S10), showed increased methylation with stDR that was associated with a decrease in Cd2 mRNA levels.

Although using oxWGBS we were able to obtain an accurate measure of true methylated cytosines in CG sites (bisulfite sequencing cannot differentiate between mCG and hmCG) in regions around the TSS in the four genes, our study lacked the sequencing depth to accurately measure changes in methylation at single CG resolution. We were only able to measure the mCG changes in regions, with our analyses focusing around the promoter. Therefore, while our study shows that stDR alters DNA methylation in the cis-acting regions of the genes studied in Lgr5^+^ ISCs, future experiments will need to critically evaluate the impact of stDNA at single CG sites. After the accurate determination of methylation changes at single CG sites, elegant experiments can be designed to modify CG sites e.g., methylate or demethylate specific CG sites or multiple CG sites to conclusively and directly test the effect of CG methylation on gene expression and ISC function. Further, the downstream effects of the genes identified to change with age and stDR (Slc28a3, Ly9, Cd2, Platr4) has to be determined to define the exact roles of these genes in age-related ISC function.

In conclusion, our study shows for the *first time* that late life stDR can restore the ability of Lgr5^+^ ISCs from old mice to proliferate and differentiate into enteroids. This observation is particularly important because it indicates that late-life strategies could be employed to rejuvenate ISCs and therefore restore gut function that declines with age and ameliorate age-related intestinal disorders. We also show for the first time that age and stDR have a major impact on the transcriptome of Lgr5^+^ ISCs and that DNA methylation potentially could mediate stDR’s effect on the ISC transcriptome especially in the old mice. Although several studies have shown that DR can impact DNA methylation in various tissues such as liver (Hahn et al., 2017; Cole et al., 2017), brain (Hadad et al., 2018) and kidney (Kim et al., 2016a), this is the first study to show that DR alters DNA methylation in ISCs. This study paves way for more research to determine the cause- and effect of DNA methylation on gene expression in ISCs.

## Methods

### Animals

Male C57BL/6 mice were obtained from the NIA aging rodent colony (Charles River Laboratories, Wilmington, MA) at 1-and 20-months of age and housed in the animal facility at the University of Oklahoma Health Sciences Center. All animals were maintained under SPF conditions in a HEPA barrier environment. The animals were fed irradiated NIH-31 mouse/rat diet from Teklad (Envigo, Madison, WI) for the entire duration of the study. At 2-and 21-months of age the mice were separated into two dietary regimens. The *ad-libitum* (AL) group and the short-term DR (stDR) group. The food consumption of all the mice in both the groups were followed for two weeks to make sure the average food consumed is same between the groups. Then the food consumed by the AL group was measured weekly for the entire duration of the study. The amount of NIH-31 diet fed each day to the DR group was adjusted based on the food consumption data of the AL group. The DR group was fed 60% of the food consumed by the mice fed AL to achieve a 40% restriction for 4-months. At the end of 4-months, the mice were sacrificed and Lgr5^+^ ISCs were isolated. The DR mice were fed at 6:00pm just before the start of light cycle. All our DR mice finish eating the food added within 1-hour of feeding. The mice were euthanized at 9:00am on the day of ISC harvesting. To match and to avoid acute effects of the 15-hour fasting experienced by our DR mice we fasted our young and old AL mice for 15-hours before the sacrifice.

### Isolation of ISCs

The isolation of Lgr5^+^ ISCs was done as previously described by Tabrizian et al., 2017. Briefly, small intestinal crypts from the mice were harvested, flushed with PBS, lumen mucosa scraped to remove, and the crypt base were mechanically dissociated to obtain crypts. Further the crypts were incubated in DMEM with 2% sorbitol for 45 min at 37° C, and the dissociated cells were collected through a cell strainer with a pore size of 10μm. Lgr5^+^ ISCs were isolated using anti-Lgr5^+^ antibody conjugated to APC and microbeads (Miltenyi, Gaithersburg, MD, USA) and sorted using column based magnetic bead sorting (Miltenyi, Gaithersburg, MD, USA). The microbeads effectively safeguard against non-specific targeting, epitope blocking, and cell activation that helps maintain target cell integrity and characteristics. Negative selection against CD45^+^ cells was done to exclude lymphocytes & a viability dye used to exclude dead cells. Similarly, paneth cells were isolated using anti-CD24^+^ antibody conjugated to microbeads.

### Real Time PCR

The levels of mRNA transcripts of ISC and paneth cell markers were measured by real-time PCR in the cells isolated using anti-Lgr5^+^ and anti-CD24^+^ antibodies conjugated with microbeads. Briefly, RNA was isolated using the RNeasy kit from Qiagen (Germantown MD, USA). The first strand cDNA was synthesized from 1µg RNA using random primers (Promega, Madison, WI, USA) and purified using the QIAquick PCR purification kit (Qiagen, Germantown, MD, USA). Expression of ISC markers, Lgr5^+^ (Mm00438890_m1), Olfm4 (Mm01320260_m1), Ascl2 (Mm01268891_g1), and Paneth Cell markers Defa22 (Mm04206099_gH), Lyz1 (Mm00657323_m1), were measured using Taqman probes and normalized to β-Actin (Mm02619580_g1). Relative gene expression was quantified as comparative ct analysis using the 2^-ΔΔct^ analysis method with β-actin as endogenous control.

### Assay of ISC function/Enteroid formation

Enteroid formation was performed as previously described by Tabrizian et al., 2017. Briefly, crypts or Lgr5^+^ ISCs isolated from the small intestine of young and old mice fed AL or stDR diet (*n* = 6/group) were washed with ADF medium, centrifuged at 300*g* for 5 min, resuspended in ADF medium and counted on a hemocytometer. Approximately 250 crypts or 8500 Lgr5^+^ ISCs were then resuspended in 25 μL of matrigel (Corning, Bedford, MA, USA), transferred to a 24-well plate to solidify at 37°C for 30 min and overlaid with 750 μl IntestiCult Organoid Growth Medium (StemCell Technologies, Vancouver, BC, Canada) and maintained at 37°C. Fresh medium was applied every 3 days and the number of total enteroids and enteroids with budding crypts were followed for 15 days and data shown for day 5 for enteroids formed from crypts and day 15 for enteroids formed from ISCs. Enteroid counting and imaging was done using inverted brightfield cell imaging system (ZOE, Bio-Rad). One-way Anova was used to determine fold difference (*p* < 0.05) with Tukey’s multiple testing correction.

For the passaging study in the old AL mice, enteroids formed from Lgr5^+^ ISCs isolated from young and old AL mice were passaged as described by Nalapareddy et al., 2017. Briefly, the medium was removed and the matrigel was dissolved in ice-cold DMEM, and then the new medium with the dissolved matrigel was pipetted 25 times with a 200-µL pipette tip. Enteroids were disrupted by passage through a 26G needle five times and re-plated in complete matrigel. Enteroids formed were counted and imaged as described above.

### Analysis of Transcriptome (RNA-Seq)

We collected transcriptomic data from two sets of samples: (i) freshly isolated total Lgr5^+^ ISCs (for Figure 3) and (ii) enteroids (for Supplement 3) by RNA-Seq as we have previously described (Unnikrishnan et al., 2017) using the Next-Generation-Sequencing Core facility at the Oklahoma Medical Research Foundation. RNA was isolated from (i) freshly isolated Lgr5^+^ ISCs and, (ii) multilobed enteroids collected from culture on day 15. Only multilobed enteroids were collected and RNA was isolated from equal number of enteroids from each group. Briefly, RNA was isolated from all 4-groups (n=6/group) using the SMART-Seq v4 Ultra Low Input RNA Kit per the manufacturer’s instructions for cDNA synthesis and amplified. Amplified cDNA was purified using Agencourt AMPure XP bead kit. Sequencing libraries were prepared using the NexteraXT DNA Library Preparation kit (Illumina FC-131-1096) with IDT-ILMN Nextera DNA UD Indexes (96 Indexes). Each library was uniquely indexed and then sized and quantified by capillary electrophoresis (TapeStation, Agilent). The indexed libraries were normalized, pooled and then sequenced on an Illumina NovaSeq 6000 system using the S1 flow cell with PE 50 reads. The raw fastq files were then imported into Strand NGS software (Version 4.0, Strand Life Sciences, Bangalore, India) for quality control, alignment, and statistical analysis. Sequences with a Q score of <30 was discarded, and the adaptor sequences were removed. Reads were aligned to mouse, build mm10 (UCSC) in an orientation-specific fashion after removing the duplicate reads. Data were normalized, and quantification was performed using the DESeq algorithm (Anders and Huber, 2010). One-way ANOVA was applied to identify differentially expressed genes with a fold change of ≥|2.0| and p < 0.05, with the Benjamini-Hochberg correction used to adjust for multiple testing. Functional annotation of candidate genes was assessed using Gene Set Enrichment Analysis of Gene Ontology (gseGO) terms with the clusterProfiler R package. Age-changes were defined as all differences found between AL-young and AL-Old animals, while diet-changes were defined as all differences between AL-young and stDR-young as well as between AL-old and stDR-old animals.

### Oxidative Whole Genome Bisulfite Sequencing (oxWGBS)

Whole Genome DNA methylation was measured using oxidative bisulfite conversion assay as previously described (Ocañas et al., 2022., Hadad et al., 2019; Hadad et al., 2016; Booth et al., 2013). To get the true 5mCG value we used oxWGBS. The traditional bisulfite sequencing technology converts unmodified cytosines to uracil to be called as unmethylated cytosine but cannot differentiate between 5mC and 5hmC and calls both as methylated cytosine. On the other hand, oxWGBS converts both the unmodified and hydroxymethyl cytosines to uracil allowing for the accurate identification of true 5mC levels. Briefly, DNA was extracted from freshly isolated Lgr5^+^ ISCs using the Qiagen DNeasy kit. Subsequently, whole genome oxidative bisulfite sequencing libraries were prepared in accordance with the manufacturer’s instructions, utilizing the Ovation Ultralow Methyl-Seq Library System from Tecan Genomics, Inc. (Redwood City, CA). About 100-300ng of genomic DNA was in a 50 µl volume of 1X low-EDTA TE buffer was used. DNA was sheared using a Covaris E220 sonicator (Covaris, Inc., Woburn, MA), and fragmented to an average fragment size of ∼200 base pairs. The quality and sizing of the sheared DNA fragments were verified using capillary electrophoresis (DNA HSD1000, Agilent), and purified using an Agencourt bead-based protocol. The purified DNA fragment in solution (100ng/12µl) were supplemented with 1 µl of CEGX spike-in control DNA (0.08 ng/µl). The subsequent steps included end repair, ligation of methylated adaptors (#L2V11DR-BC 1–96 adaptor plate, Tecan Genomics), and final repair, performed as per the manufacturer’s instructions. The resulting DNA was subjected to oxidation and bisulfite conversion using the True Methyl oxBS module (NuGEN), including desulfonation and purification steps. Following purification, 22 µl of libraries were eluted using magnetic beads. Oxidative bisulfite-converted samples were subsequently amplified for 15 cycles [95°C-2 min, N (95°C-15 s, 60°C-1 min, 72°C-30 s]. The resulting amplified libraries were purified using Agencourt beads and eluted in low-EDTA TE buffer. Validation and quantification of the libraries were performed using capillary electrophoresis (TapeStation HSD1000, Agilent). The amplified libraries were then normalized to a concentration of 4nM, pooled, and subjected to denaturation and dilution, resulting in a final concentration of 12pM. Finally, a custom sequencing primer, MetSeq Primer, was introduced and spiked in with the Illumina Read 1 primer to achieve a final concentration of 0.5 µM. Libraries were then sequenced on an Illumina Novaseq 6000 with PE150bp reads.

Data analyses were done as previously described (Chucair-Elliott et al., 2020). Briefly, quality checks on fastq files were performed using FastQC (https://www.bioinformatics.babraham.ac.uk/projects/fastqc/). Paired-end reads then underwent preprocessing with Trimmomatic version 0.39 (Bolger AM et al., 2014), where adaptor trimming was performed. Leading and trailing bases with Q-scores below 25 were removed, reads shorter than 30 bases were discarded, and reads with an average Q-score below 25 were filtered out. Trimmed reads were then aligned to the mouse reference genome (GRCm38/mm10) using Bismark (Felix K et al., 2011) with Bowtie2 (Langmead B et al., 2012). Resulted Bams were deduplicated using Bismark and sorted using SAMtools sort (Heng Li et al., 2009). Sorted BAM files were then run through MethylKit in R (Akalin et al. 2012) to generate context-specific (CG/CH) coverage text files.

We then established flanking regions spanning 4000 nucleotides (including shores and shelves), both upstream of each gene’s transcription start site and downstream of the gene’s end site for *Slc28a3, Ly9, Cd2, and Platr4*. These regions were partitioned into 20 bins, each spanning 200 nucleotides and the gene bodies were divided into 27 equal bins, with the bin count adjusted according to the length of the gene, to facilitate granular analysis. We subsequently determined average mCG levels around the transcription start site (TSS) by focusing on a 1000-base pair span, partitioned into 10 bins of 100 nucleotides each, encompassing both the upstream region (500 bases) and the downstream region (500 bases) around the TSS. Age-changes were defined as all differences found between AL-young and AL-Old animals, while diet-changes were defined as all differences between AL-young and stDR-young as well as between AL-old and stDR-old animals.

### Statistical Analysis

All data are summarized as mean ± SEM. Statistical tool used, and the n (number of mice) used for each experiment is mentioned in the respective figure legend. One-way ANOVA (main effect) followed by multiple testing corrections (Tukey’s or Benjamini-Hochberg) was used as the statistical tool to determine significance. To illustrate significance, we used GraphPad version 10.0.2 for Windows, GraphPad Software, Boston, Massachusetts USA, www.graphpad.com.

## Acknowledgements

The research was supported by the following grants: NIH grants P20GM125528 Geroscience CoBRE (AU), KO1AG 056655-01A1 (AU), and Oklahoma Center for Adult Stem Cell Research (AU), VA grants IK6BX006033 (WMF), I01BX003906 (WMF) and NIH grants P30AG050911 (WMF), R01AG059430 (WMF).

