## Supplement for "Late-life short-term dietary restriction ameliorates intestinal stem cell function and impacts intestinal stem cell DNA methylation": Supplement_KurupChan et al 2024.pptx

### Slide 1
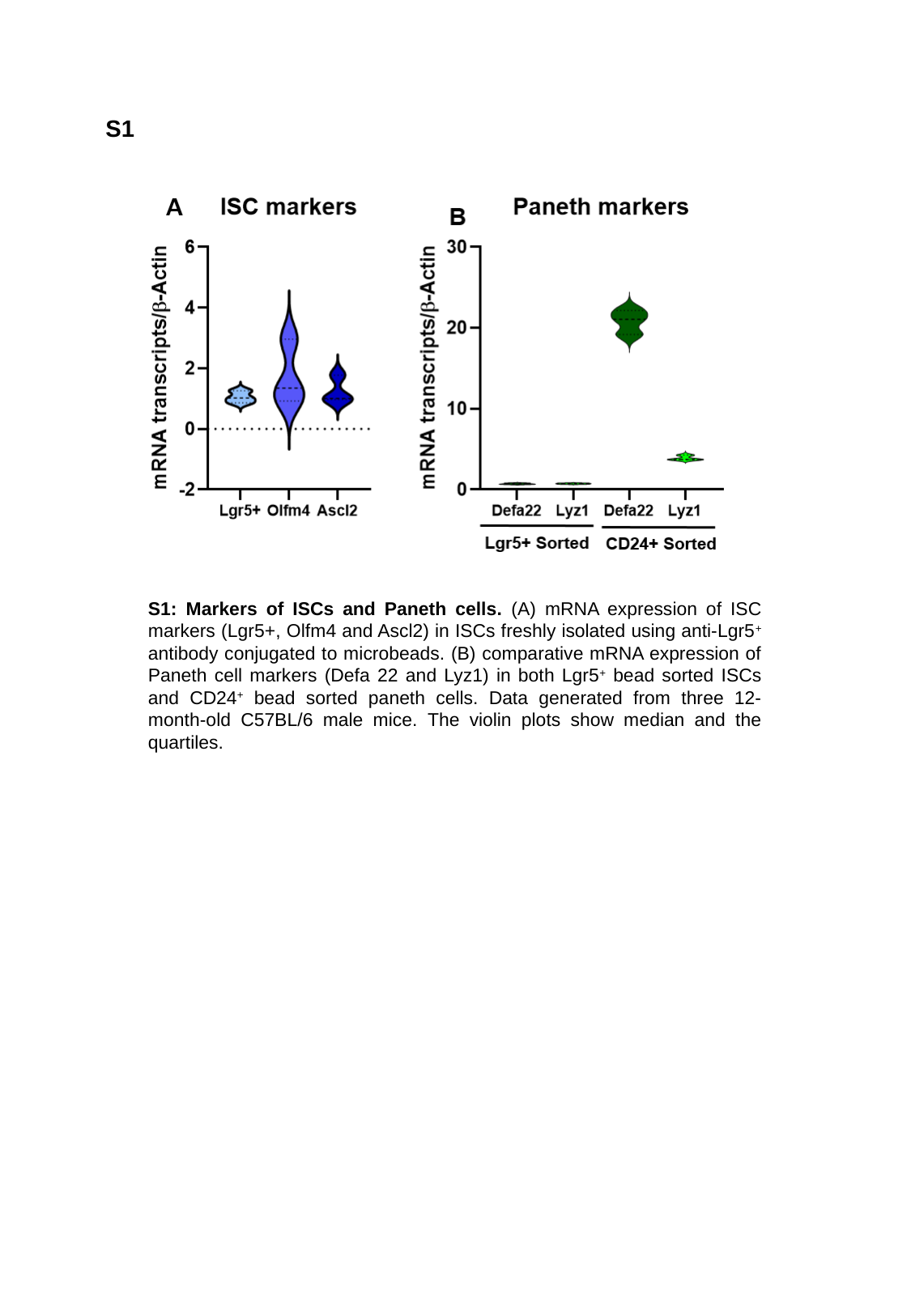

S1
Young
S1: Markers of ISCs and Paneth cells. (A) mRNA expression of ISC markers (Lgr5+, Olfm4 and Ascl2) in ISCs freshly isolated using anti-Lgr5^+^ antibody conjugated to microbeads. (B) comparative mRNA expression of Paneth cell markers (Defa 22 and Lyz1) in both Lgr5^+^ bead sorted ISCs and CD24^+^ bead sorted paneth cells. Data generated from three 12-month-old C57BL/6 male mice. The violin plots show median and the quartiles.

### Slide 2
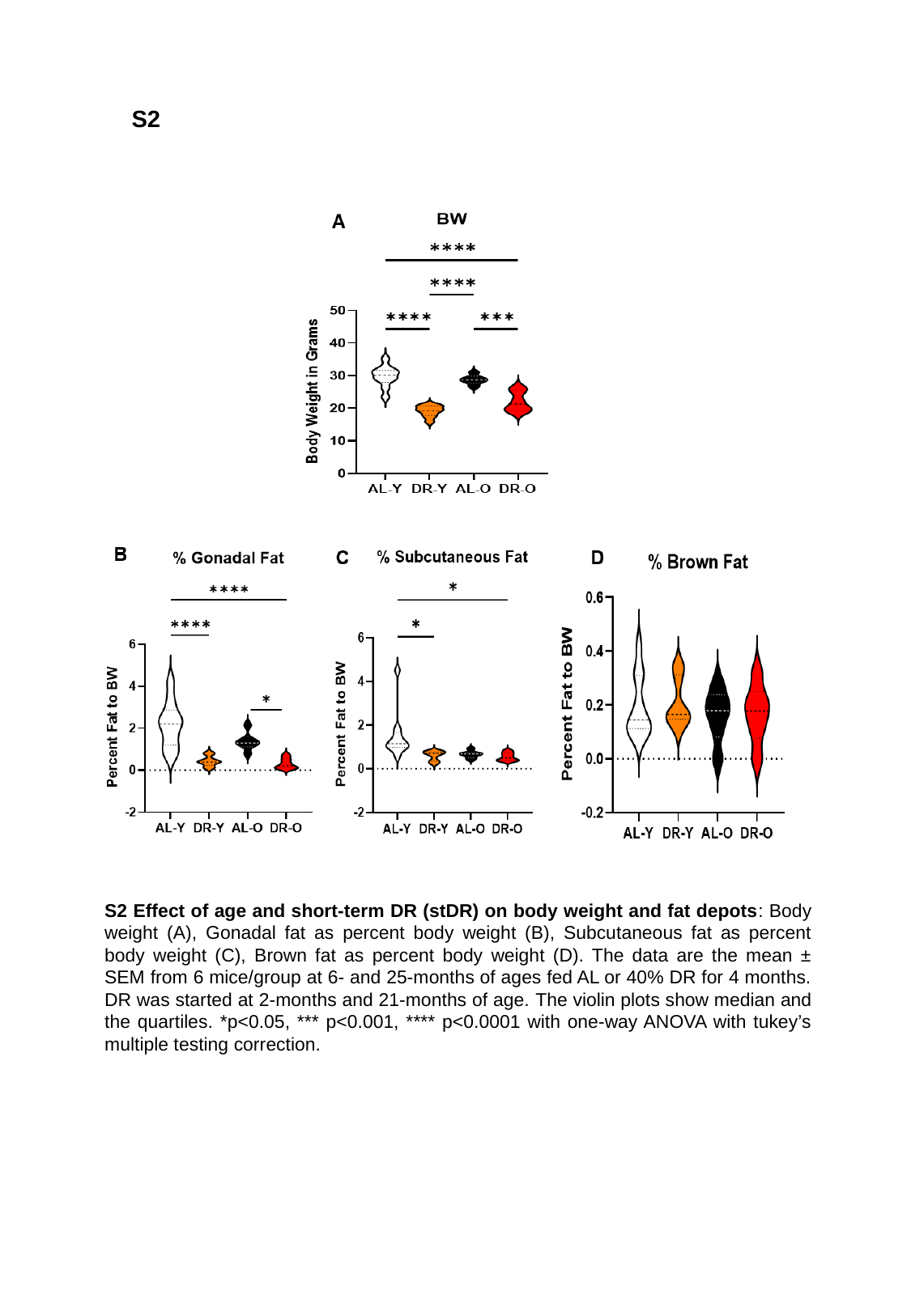

S2
S2 Effect of age and short-term DR (stDR) on body weight and fat depots: Body weight (A), Gonadal fat as percent body weight (B), Subcutaneous fat as percent body weight (C), Brown fat as percent body weight (D). The data are the mean ± SEM from 6 mice/group at 6- and 25-months of ages fed AL or 40% DR for 4 months. DR was started at 2-months and 21-months of age. The violin plots show median and the quartiles. *p<0.05, *** p<0.001, **** p<0.0001 with one-way ANOVA with tukey’s multiple testing correction.

### Slide 3
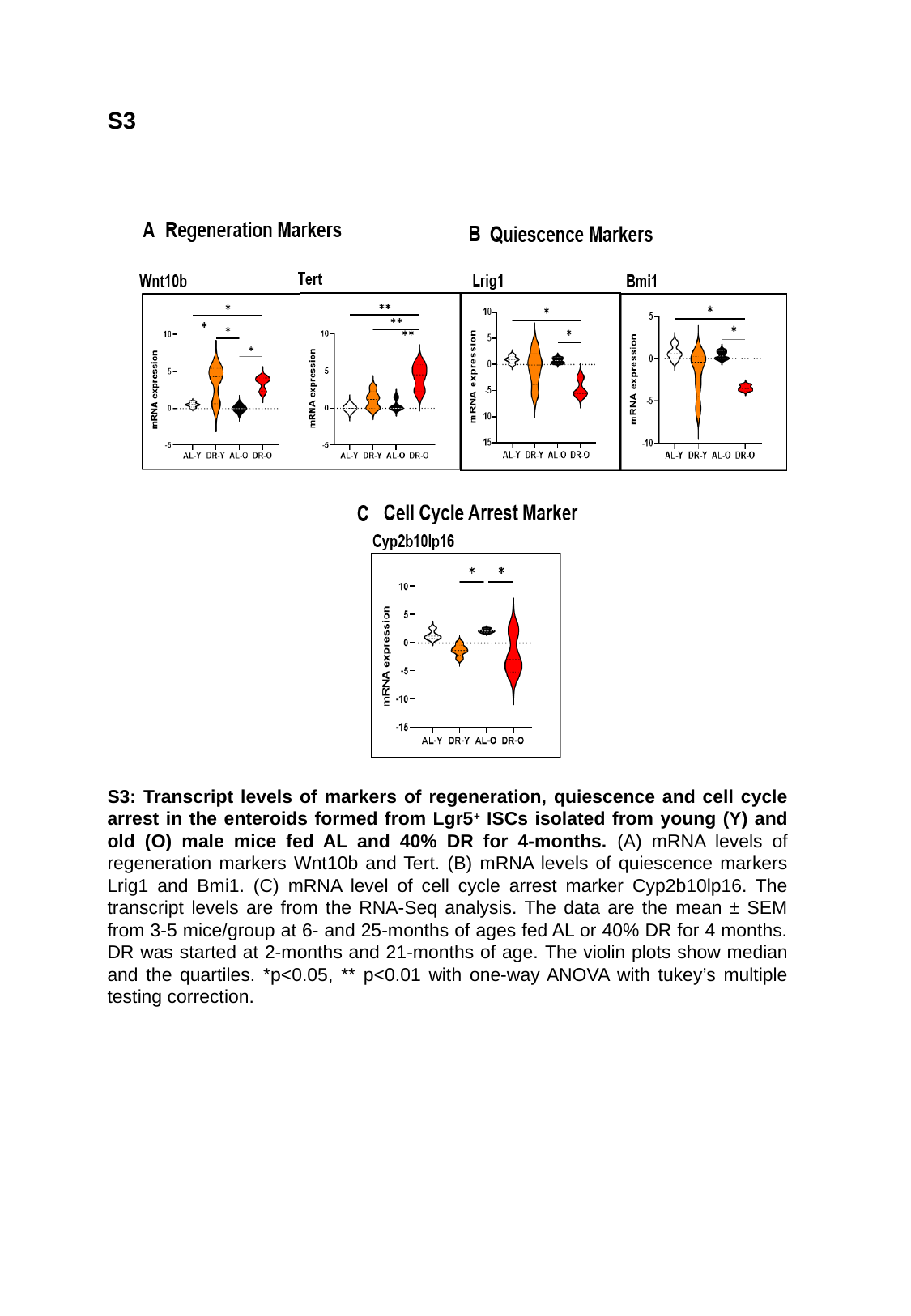

S3
S3: Transcript levels of markers of regeneration, quiescence and cell cycle arrest in the enteroids formed from Lgr5^+^ ISCs isolated from young (Y) and old (O) male mice fed AL and 40% DR for 4-months. (A) mRNA levels of regeneration markers Wnt10b and Tert. (B) mRNA levels of quiescence markers Lrig1 and Bmi1. (C) mRNA level of cell cycle arrest marker Cyp2b10lp16. The transcript levels are from the RNA-Seq analysis. The data are the mean ± SEM from 3-5 mice/group at 6- and 25-months of ages fed AL or 40% DR for 4 months. DR was started at 2-months and 21-months of age. The violin plots show median and the quartiles. *p<0.05, ** p<0.01 with one-way ANOVA with tukey’s multiple testing correction.

### Slide 4
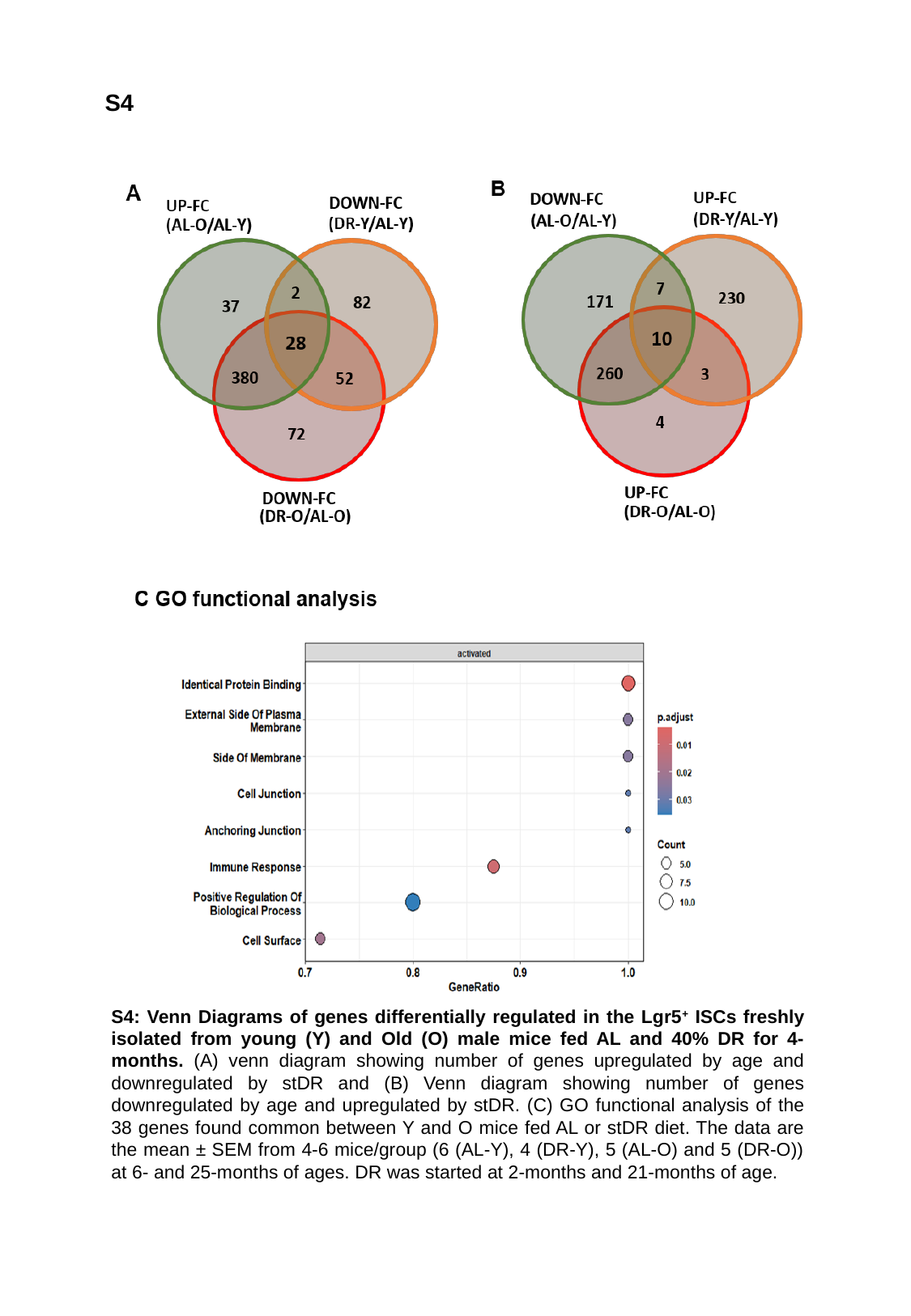

S4
S4: Venn Diagrams of genes differentially regulated in the Lgr5^+^ ISCs freshly isolated from young (Y) and Old (O) male mice fed AL and 40% DR for 4-months. (A) venn diagram showing number of genes upregulated by age and downregulated by stDR and (B) Venn diagram showing number of genes downregulated by age and upregulated by stDR. (C) GO functional analysis of the 38 genes found common between Y and O mice fed AL or stDR diet. The data are the mean ± SEM from 4-6 mice/group (6 (AL-Y), 4 (DR-Y), 5 (AL-O) and 5 (DR-O)) at 6- and 25-months of ages. DR was started at 2-months and 21-months of age.

### Slide 5
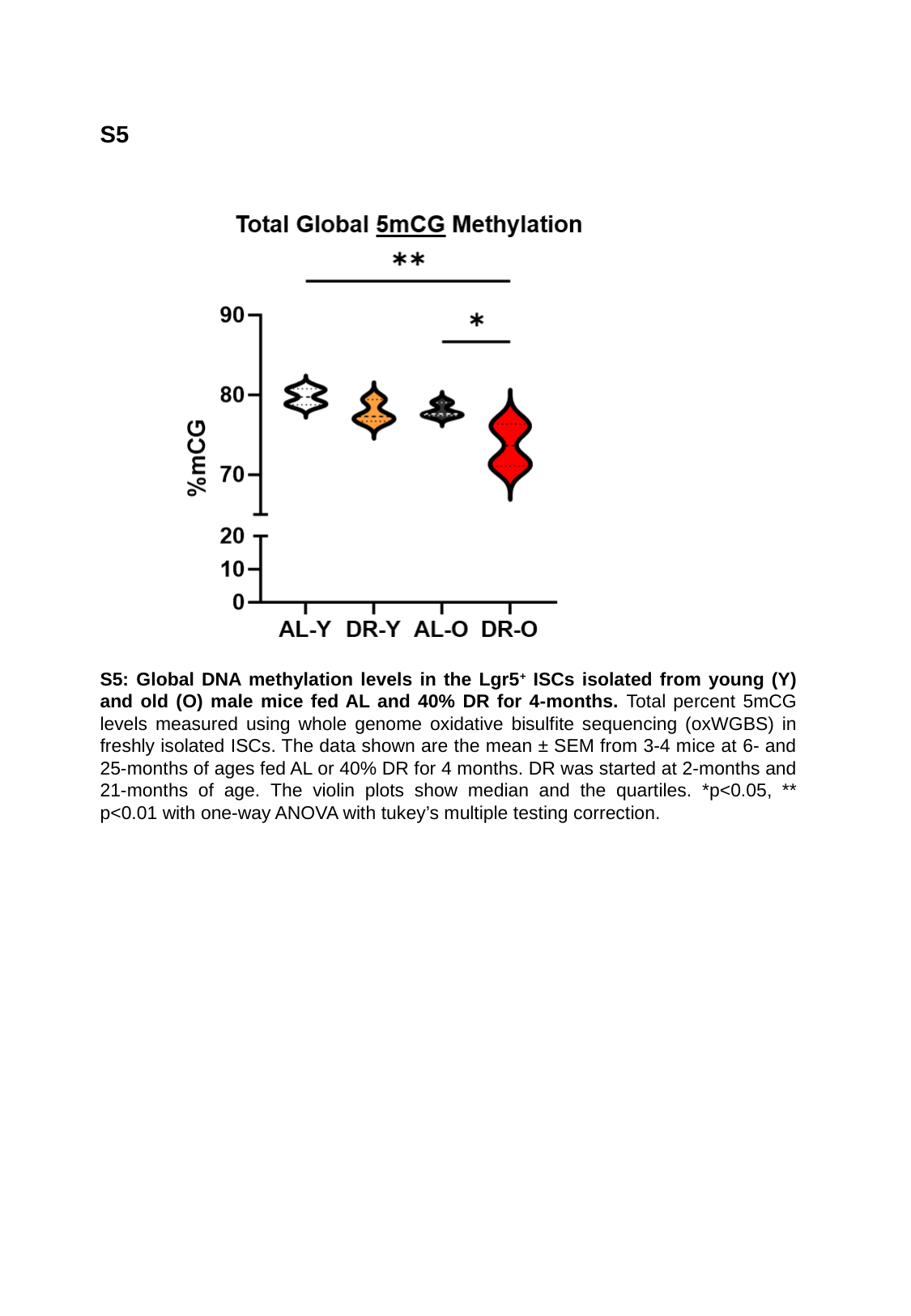

S5
S5: Global DNA methylation levels in the Lgr5^+^ ISCs isolated from young (Y) and old (O) male mice fed AL and 40% DR for 4-months. Total percent 5mCG levels measured using whole genome oxidative bisulfite sequencing (oxWGBS) in freshly isolated ISCs. The data shown are the mean ± SEM from 3-4 mice at 6- and 25-months of ages fed AL or 40% DR for 4 months. DR was started at 2-months and 21-months of age. The violin plots show median and the quartiles. *p<0.05, ** p<0.01 with one-way ANOVA with tukey’s multiple testing correction.

### Slide 6
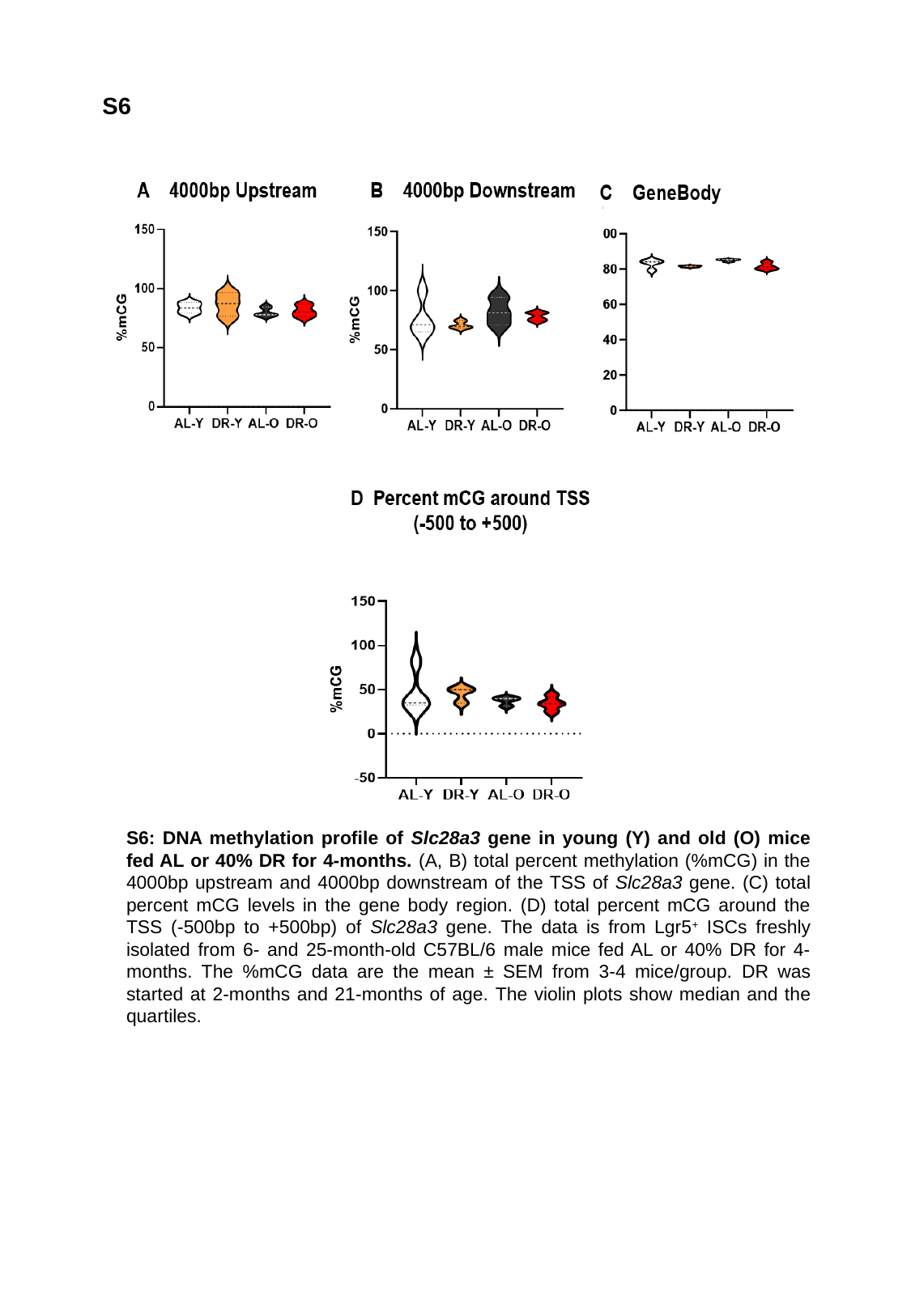

S6
S6: DNA methylation profile of Slc28a3 gene in young (Y) and old (O) mice fed AL or 40% DR for 4-months. (A, B) total percent methylation (%mCG) in the 4000bp upstream and 4000bp downstream of the TSS of Slc28a3 gene. (C) total percent mCG levels in the gene body region. (D) total percent mCG around the TSS (-500bp to +500bp) of Slc28a3 gene. The data is from Lgr5^+^ ISCs freshly isolated from 6- and 25-month-old C57BL/6 male mice fed AL or 40% DR for 4-months. The %mCG data are the mean ± SEM from 3-4 mice/group. DR was started at 2-months and 21-months of age. The violin plots show median and the quartiles.

### Slide 7
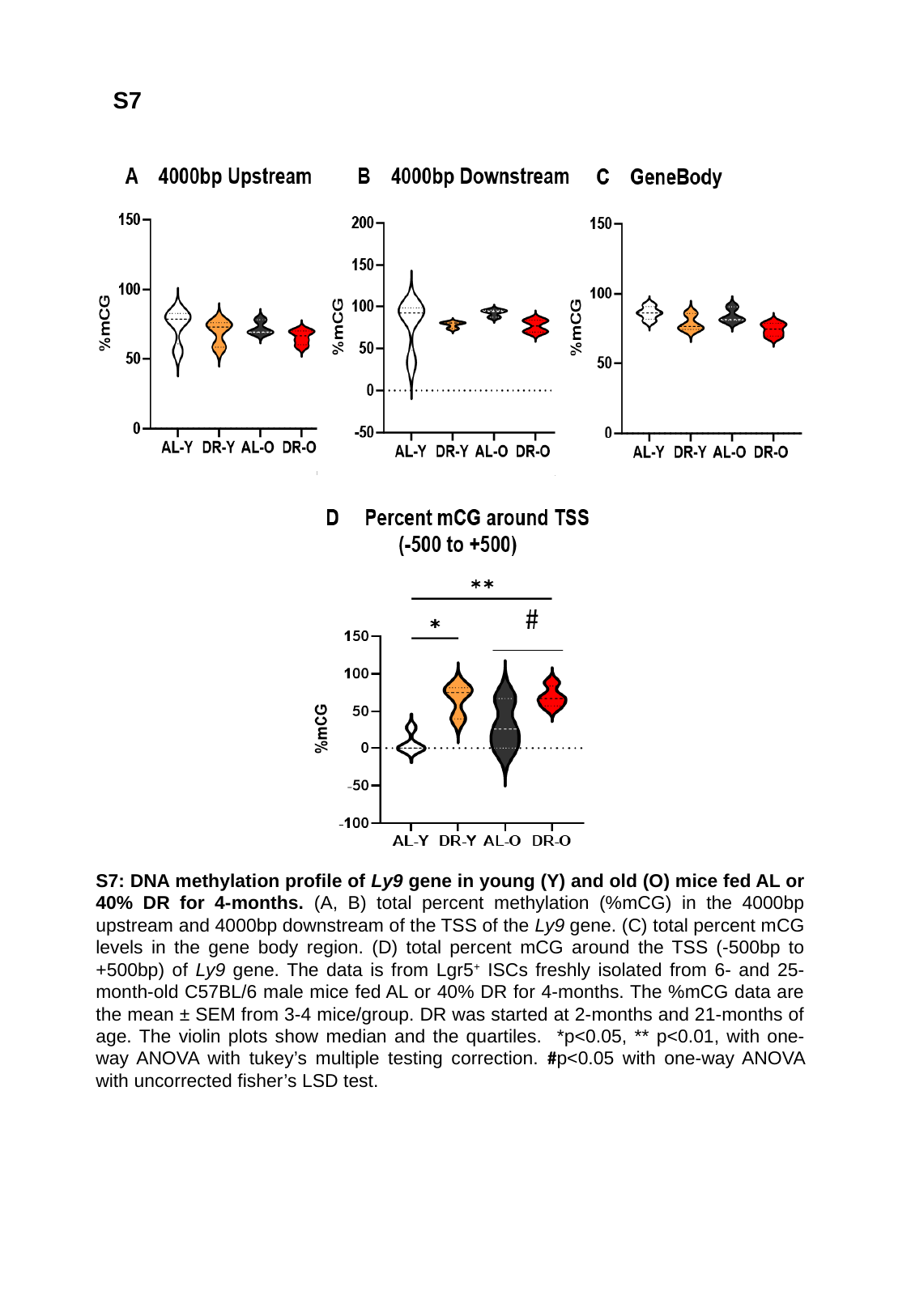

S7
S7: DNA methylation profile of Ly9 gene in young (Y) and old (O) mice fed AL or 40% DR for 4-months. (A, B) total percent methylation (%mCG) in the 4000bp upstream and 4000bp downstream of the TSS of the Ly9 gene. (C) total percent mCG levels in the gene body region. (D) total percent mCG around the TSS (-500bp to +500bp) of Ly9 gene. The data is from Lgr5^+^ ISCs freshly isolated from 6- and 25-month-old C57BL/6 male mice fed AL or 40% DR for 4-months. The %mCG data are the mean ± SEM from 3-4 mice/group. DR was started at 2-months and 21-months of age. The violin plots show median and the quartiles. *p<0.05, ** p<0.01, with one-way ANOVA with tukey’s multiple testing correction. #p<0.05 with one-way ANOVA with uncorrected fisher’s LSD test.

### Slide 8
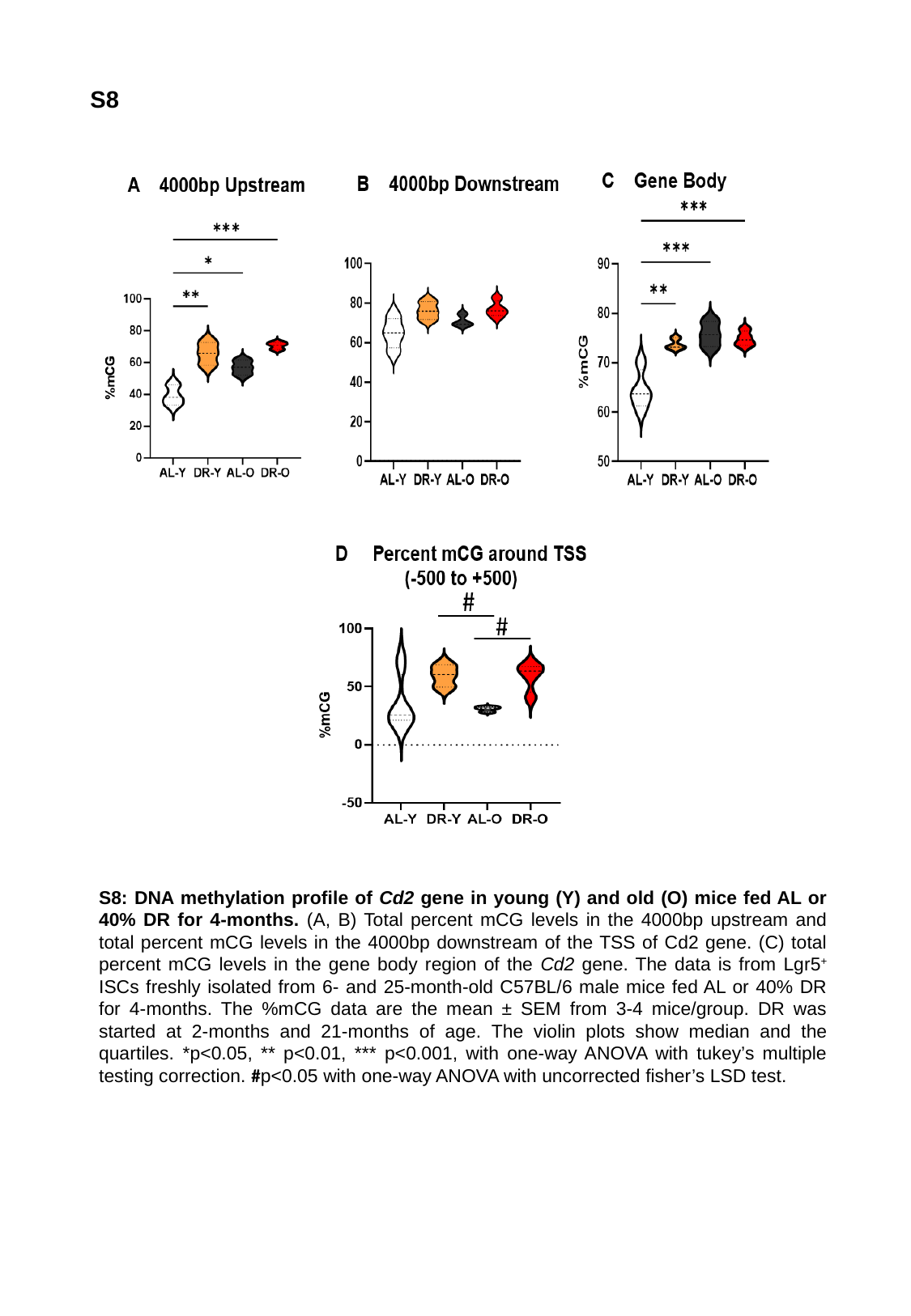

S8
S8: DNA methylation profile of Cd2 gene in young (Y) and old (O) mice fed AL or 40% DR for 4-months. (A, B) Total percent mCG levels in the 4000bp upstream and total percent mCG levels in the 4000bp downstream of the TSS of Cd2 gene. (C) total percent mCG levels in the gene body region of the Cd2 gene. The data is from Lgr5^+^ ISCs freshly isolated from 6- and 25-month-old C57BL/6 male mice fed AL or 40% DR for 4-months. The %mCG data are the mean ± SEM from 3-4 mice/group. DR was started at 2-months and 21-months of age. The violin plots show median and the quartiles. *p<0.05, ** p<0.01, *** p<0.001, with one-way ANOVA with tukey’s multiple testing correction. #p<0.05 with one-way ANOVA with uncorrected fisher’s LSD test.

### Slide 9
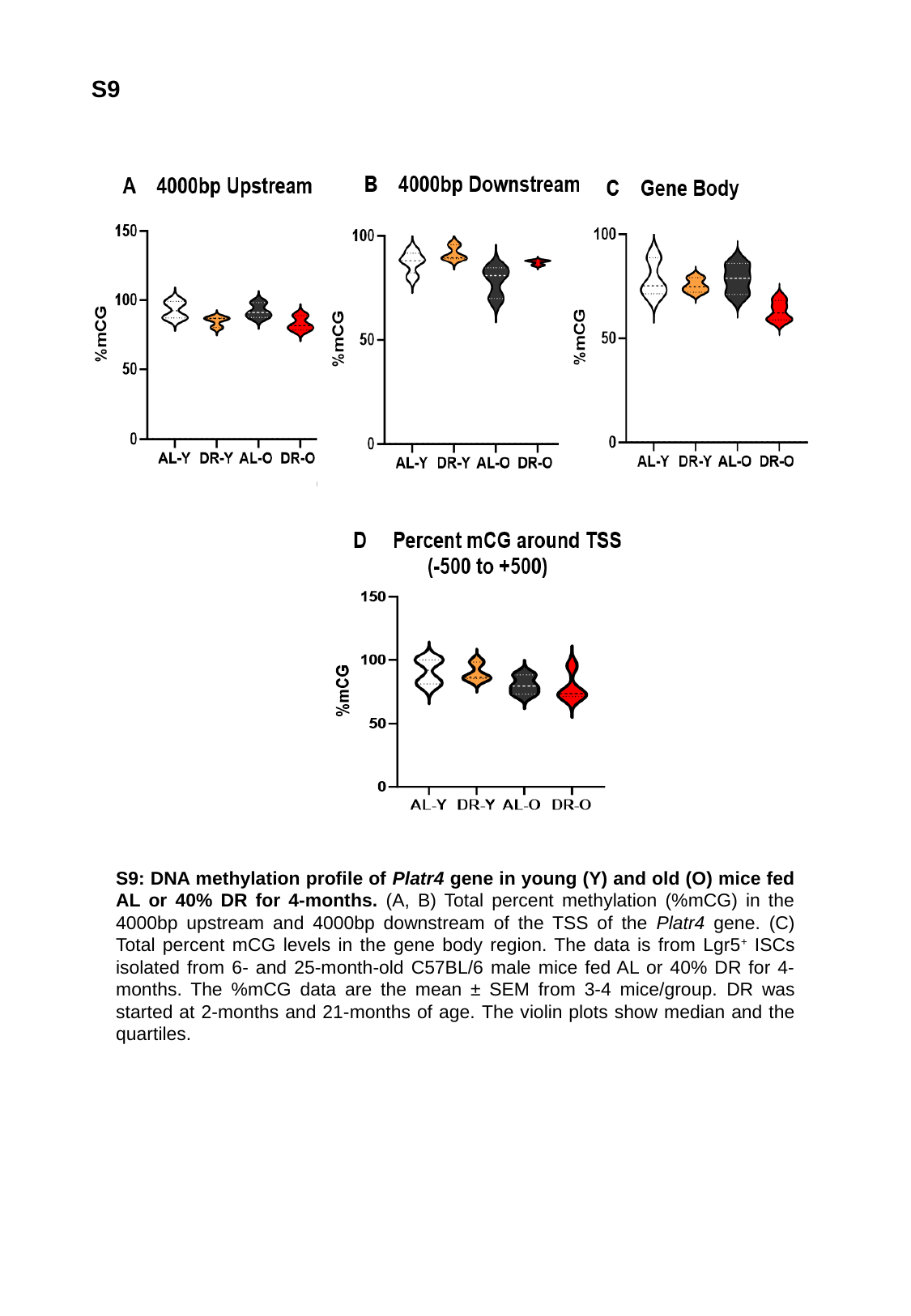

S9
S9: DNA methylation profile of Platr4 gene in young (Y) and old (O) mice fed AL or 40% DR for 4-months. (A, B) Total percent methylation (%mCG) in the 4000bp upstream and 4000bp downstream of the TSS of the Platr4 gene. (C) Total percent mCG levels in the gene body region. The data is from Lgr5^+^ ISCs isolated from 6- and 25-month-old C57BL/6 male mice fed AL or 40% DR for 4-months. The %mCG data are the mean ± SEM from 3-4 mice/group. DR was started at 2-months and 21-months of age. The violin plots show median and the quartiles.

### Slide 10
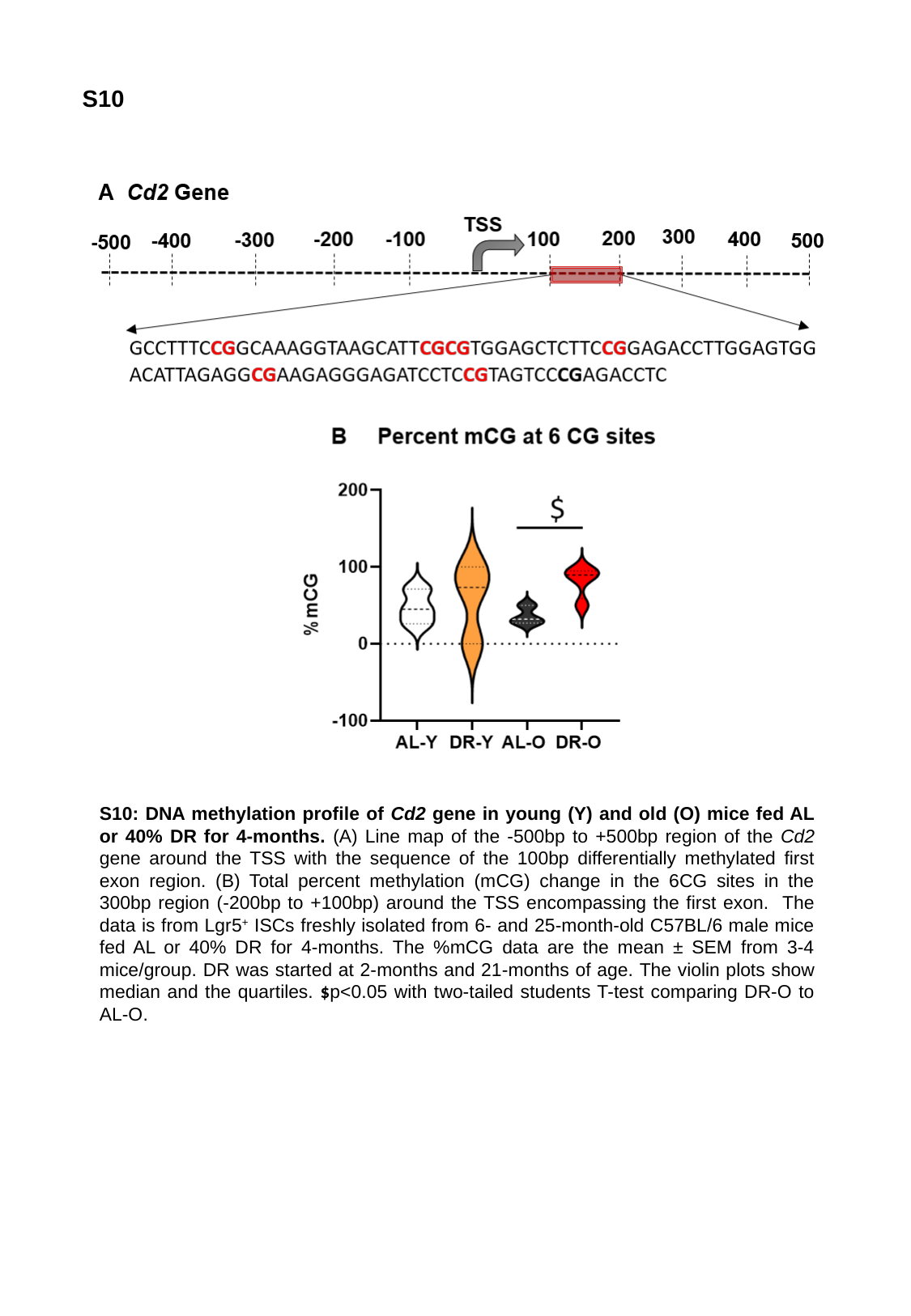

S10
S10: DNA methylation profile of Cd2 gene in young (Y) and old (O) mice fed AL or 40% DR for 4-months. (A) Line map of the -500bp to +500bp region of the Cd2 gene around the TSS with the sequence of the 100bp differentially methylated first exon region. (B) Total percent methylation (mCG) change in the 6CG sites in the 300bp region (-200bp to +100bp) around the TSS encompassing the first exon. The data is from Lgr5^+^ ISCs freshly isolated from 6- and 25-month-old C57BL/6 male mice fed AL or 40% DR for 4-months. The %mCG data are the mean ± SEM from 3-4 mice/group. DR was started at 2-months and 21-months of age. The violin plots show median and the quartiles. $p<0.05 with two-tailed students T-test comparing DR-O to AL-O.
